# An exploratory computational analysis in mice brain networks of widespread epileptic seizure onset locations along with potential strategies for effective intervention and propagation control

**DOI:** 10.1101/2023.12.22.573015

**Authors:** Juliette Courson, Mathias Quoy, Yulia Timofeeva, Thanos Manos

## Abstract

Mean-field models have been developed to replicate key features of epileptic seizure dynamics. However, the precise mechanisms and the role of the brain area responsible for seizure onset and propagation remain incompletely understood. In this study, we employ computational methods within The Virtual Brain framework and the Epileptor model to explore how the location and connectivity of an Epileptogenic Zone (EZ) in a mouse brain are related to focal seizures (seizures that start in one brain area and may or may not remain localized), with a specific focus on the hippocampal region known for its association with epileptic seizures. We then devise computational strategies to confine seizures (prevent widespread propagation), simulating medical-like treatments such as tissue resection and the application of an anti-seizure drugs or neurostimulation to suppress hyperexcitability. Through selectively removing (blocking) specific connections informed by the structural connectome and graph network measurements or by locally reducing outgoing connection weights of EZ areas, we demonstrate that seizures can be kept constrained around the EZ region. We successfully identified the minimal connections necessary to prevent widespread seizures, with a particular focus on minimizing surgical or medical intervention while simultaneously preserving the original structural connectivity and maximizing brain functionality.

## 1 INTRODUCTION

An essential objective of computational neuroscience is to be able to predict the effects of medical interventions, therapies, or surgeries on the local and global brain dynamical activity. A dynamical-model approach can establish connections among attractors, bifurcations, synchronization patterns, and empirical neuroimaging data like EEG and fMRI. By appropriately selecting model parameters it allows us to construct tailored in silico brain activity profiles for different subjects. These parameter settings can function as dynamic indicators and prognosticators of distinct brain states (healthy vs. pathological) and behavioral patterns (Popovych et al., 2019). Such an approach would enable the customization of treatments applied for example in brain stimulation therapies for different patients. In the latter ones, the clinical goal is to reverse the irregular neural synchronization linked to various neurological disorders such as Parkinson’s disease and tinnitus (Tass, 2003; Manos et al., 2018a,b, 2021). Mathematical models can provide a theoretical framework to investigate the impact of the localised population dynamics combined with the complex topology of the brain network on the overall brain activity (Izhikevich, 2007; et al., 2014) as well as to account for different types of plasticity (e.g., Hebbian or structural plasticity) and their impact on the system’s dynamics (Berry and Quoy, 2006; Pitti et al., 2017, 2020; Butz et al., 2009; Ooyen and Butz-Ostendorf, 2017; Manos et al., 2021; Bi et al., 2021; Lu et al., 2023; Bergoin et al., 2023).

Among these neurological disorders, epilepsy is a very common neurological condition that affects over 46 million people worldwide (Beghi, 2019). It is characterized by the occurrence of multiple epileptic seizures, causing a wide range of symptoms including language and motor troubles, auras, convulsions and loss of consciousness (Chang et al., 2017). These seizures are generated by either a localized production of excessive discharges occurring in one area or hemisphere (i.e. *focal*), or simultaneously in both hemispheres (i.e. *generalized*), that can either remain localized or propagate in the brain network (widespread seizure) (Englot et al., 2016; Burman and Parrish, 2018). Around 30% of the epileptic patients are drug-resistant, and therefore may have to undergo resective surgical intervention to cure their disease, which usually consists of the removal of the entire Epileptogenic Zone (EZ) (Jehi, 2018; Rubio et al., 2019). In Temporal Lobe Epilepsy, around 70% of the resective surgery operations lead to the suppression of the epileptic activity or the drastic diminution of its occurrence, but also causes brain dysfunctions, especially memory impairment. The surgery is ineffective in around 30% of the cases (Alexandratou et al., 2021). It is evident that adequately identifying the epileptogenic brain area is crucial to perform appropriate resections. Usually based on visual inspections or quantitative analysis of intracranial electroencephalogram (IEEG) data, the detection of the EZ may not succeed in cases of complex seizure initiation patterns (Andrzejak et al., 2014).

Over the past years there has been an increasing integration of neuroimaging data to enhance the accuracy and predictive capacity of mathematical models (Popovych et al., 2019, 2021; Manos et al., 2023). This integration occurs within a virtual (computational) environment where simulations enable the exploration and derivation of parameters that could eventually facilitate the prediction of micro- and macro-scale brain activity states (Shusterman and Troy, 2008; Jirsa et al., 2017; Chizhov and Sanin, 2020). Personalized models of epileptic brain using patient-specific data allow us to test various resection options before surgery (see e.g., (Jirsa et al., 2016)), and the relevance of preferentially removing connections based on their location in the modular brain structure (see e.g. (Olmi et al., 2019; Nissen et al., 2021)). In (An et al., 2019) the authors used mathematical models and brain network simulations, coupled with modularity analysis based on individual structural brain connectivity to pinpoint optimal surgical target areas. Subsequent to this investigation, in (Hashemi et al., 2020) the authors introduced a probabilistic framework capable of inferring the spatial distribution of epileptogenicity within a personalized, large-scale brain model of epilepsy propagation. In (Makhalova et al., 2022), the authors used such computational approaches to compare the regions identified as epileptogenic by the so-called Virtual epileptic patient brain model to those defined by clinical analysis (see also (Jirsa et al., 2023)).

A viable alternative is to use virtual mice models. Indeed, rodent species are often regarded as suitable analogs for humans due to the significant similarities in brain structure and connectivity between the two (Grone and Baraban, 2015; Marshall et al., 2021). Scientists frequently opt for laboratory mice as research subjects because these creatures also bear genetic resemblance to humans and have shorter lifespans, enabling multi-generational studies. Furthermore, advancements in imaging technology are rapidly enhancing the precision and detail of experimental data. The latest iterations of MRI machines, for instance, provide intricate insights into the anatomical, structural, and functional aspects of the entire rodent brain (Stafford et al., 2014). The primary objective of mouse-based research is to deepen our comprehension of brain function and malfunction. The ultimate goal is to acquire new knowledge of the mechanisms controlling and intervening in the brain dynamics which can be later utilised in human brain’s therapies, like for example in the treatment of Epilepsy.

A better understanding of the role of neural network’s topological properties in the spatiotemporal propagation of epileptic activity would open the path for more effective EZ identification, and better resection strategies. In (Toyoda et al., 2013), the authors used recording electrodes to assess the propagation of seizures in rats experiencing spontaneous seizures. They observed that the initial seizure activity was most frequently detected in the hippocampal formation, followed by sequential spreading to the subiculum, entorhinal cortex, olfactory cortex, neocortex, and striatum. In (Melozzi et al., 2017), the authors numerically simulated the dynamical behavior of a mouse brain to replicate certain aspects of the anatomical reorganization observed in medial temporal lobe epilepsy reported in (Toyoda et al., 2013). To this end, they focused on the loss of neuronal connections in the hippocampal regions (CA1 and CA3) [see also experimental findings in (Esclapez et al., 1999)] and they eliminated (in silico) all incoming and outgoing connections of CA1 and CA3 brain areas in the hippocampus to prevent widespread epileptic seizure propagation.

Epileptic seizures arise from an imbalance in the regulation of stimulation and inhibition. Cellular-level processes involve ion transporters, pumps, and channels that govern the entry and exit of positively or negatively charged ions within neurons. These mechanisms are further modulated by factors such as voltage or ligands, either binding directly or through G protein receptors, which exert control over these pumps and ion channels (see (Bakhtiarzadeh et al., 2023) and references therein). There are several anti-seizure drugs that have been reported to suppress epileptiform spikes and improve synaptic and cognitive function (see (Kanner and Bicchi, 2022) for a recent review). In parallel, neuromodulation techniques have emerged as promising strategies for influencing brain activity and have gained considerable attention in the context of managing seizure propagation. One of the key players in this field is transcranial magnetic stimulation (TMS), which non-invasively modulates neuronal excitability by generating magnetic fields that induce electrical currents in targeted brain regions. Studies have shown that repetitive TMS (rTMS) can alter cortical excitability and disrupt abnormal synchronization of neuronal networks, thereby attenuating seizure propagation, see e.g., (Tsuboyama et al., 2020) and references therein. Additionally, vagus nerve stimulation (VNS) has demonstrated effectiveness in reducing seizure frequency and severity by modulating the autonomic nervous system and releasing neurotransmitters that promote inhibitory signaling, see e.g., (Toffa et al., 2020) and references therein. Moreover, recent advancements in closed-loop neuromodulation, such as responsive neurostimulation (RNS), offer real-time monitoring and adaptive delivery of electrical pulses to preemptively suppress abnormal neuronal activity and prevent seizure spread (Heck et al., 2014). These findings highlight the potential of neuromodulation techniques as adjunctive therapies for controlling seizure propagation and improving the quality of life for individuals with epilepsy.

In this work, we computationally study how the location of an EZ area and its connectivity relevance in the network are related to widespread seizure propagation in a mice brain and we search for strategies that can confine widespread seizures by either removing the minimum amount of brain tissue (by blocking certain connections in the network), or suppress the hyperexcitation (loosely mimicking a anti-seizure drug or neuromodulation effect). To this end, instead of (computationally) resecting the whole EZ tissue from both brain hemispheres as in (Melozzi et al., 2017) (via a graph edge removal), we sought out to systematically identify and remove (or block) the minimal amount of connections required to prevent a widespread seizure propagation. In addition, we followed an alternatively approach to computationally model (in a loose sense) the effect of a drug at a macroscopic scale (i.e., suppression of network hyperexcitability). To this objective, we altered locally the outgoing weight connections in our structural connectome to account for the inhibitory effect of such a drug in the vicinity of a given EZ area. In both of our approaches, the ultimate goal is to minimize the surgical or medical intervention and preserve as much as possible the pre-surgical structural connectivity as well as the maximum possible amount of the brain functionality.

## 2 MATERIALS AND METHODS

### The Epileptor model

The Epileptor (Jirsa et al., 2014; Proix et al., 2014; Houssaini et al., 2020) is a phenomenological model for the description of local-field potentials during epileptic seizures, comprised by one susbsystem with two state variables (*x*_1_, *y*_1_) responsible for generating fast discharges Hindmarsh and Rose, 1984) and a second one with two state variables (*x*_2_, *y*_2_) generating sharp-wave events (Roy et al., 2011). The fast and slow variables are linked to the so-called permittivity variable, *z* (see below) which evolves on a very slow timescale. The onset of a seizure takes place through a saddle-node bifurcation while the time evolution and the offset via a homoclinic bifurcation. The Epileptor was also used in (Melozzi et al., 2017) to model epileptic activity for mouse brains. In our study, we consider a network of *N* Epileptors, coupled via an adjacency matrix *A* = (*c*_*ji*_) while we do not include any delays using track length data. The epileptic seizure-like events are produced by the model described with the following set of equations:

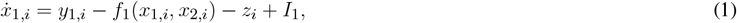

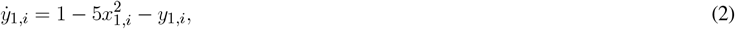

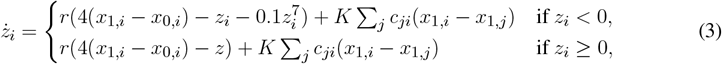

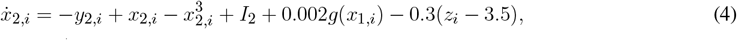

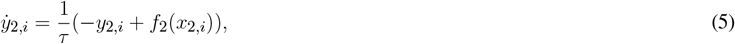

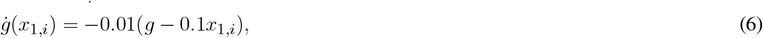

where

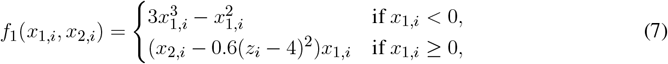

and

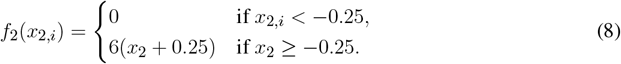

The combined *x*_2,*i*_ − *x*_1,*i*_ variable models the potential of brain area *i*. Following the work of (Melozzi et al., 2017) and tuning our system in a similar state, we set *I*_1_ = 3.1, *I*_2_ = 0.45, *r* = 3.5.10^−4^, *K* = 0.2, *τ* = 10 s. Additive Gaussian white noise with standard deviation *σ* = 0.0025 is added to Eqs. (4) and (5). Our simulations were performed with The Virtual Brain (TVB) platform Sanz Leon et al. (2013); Sanz-Leon et al. (2015).

In our work, and for each brain node, we focus only on the epileptogenicity parameter (*x*_0,*i*_) and we adjust its value according to healthy or epileptic condition of each node. The critical epileptogenicity parameter for the transition from one state to the other is found to be 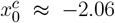 (see (Houssaini et al., 2020)). A node or group of EZ nodes (Jehi, 2018) is selected to produce spontaneous epileptic seizures, while the rest of the brain nodes form the healthy Propagation Zone (PZ). We set the values of the epileptogenicity in these brain nodes as *x*_0_ = *−*1.6 for the EZ and *x*_0_ = *−*2.1 for the PZ nodes.

In order to systematically detect the seizure onset for each brain area and its widespread (or not) evolution to other regions, we use the evolution of the slow permittivity *z*. First, we allow the system to evolve for approximately 15 seconds (this also depends on the onset of a first seizure) to eliminate temporary effects. At this stage all PZ nodes are in a stable state (i.e., the *z* variable value is approximately constant), and the EZ does not produce a seizure yet. Therefore, we define seizure onset time *T*_onset,*i*_ in each brain area *i*, as the time when the slow permittivity (*z*) starts increasing. To track the propagation of the seizure among brain areas, we define the time distance from seizure onset in each network node *i*:

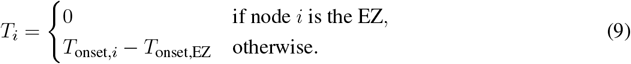

with *T*_onset,EZ_ the onset time of the seizure in the EZ.

In **Figure** 1, we show examples of an epileptogenic (in red) and healthy (in blue) nodes activity produced by the Epileptor model. The time evolution of the fast variable *x*_2_*− x*_1_ is depicted in blue (non EZ) and red (initially an EZ). The slow permittivity variable *z* is in black. Non-epileptogenic brain areas can either be recruited in the seizure, or maintain a healthy activity. Some examples detecting the time stamps of seizure onset in each node using the slow variable *z* are also depicted in **Figure** 1.

**Figure 1.**
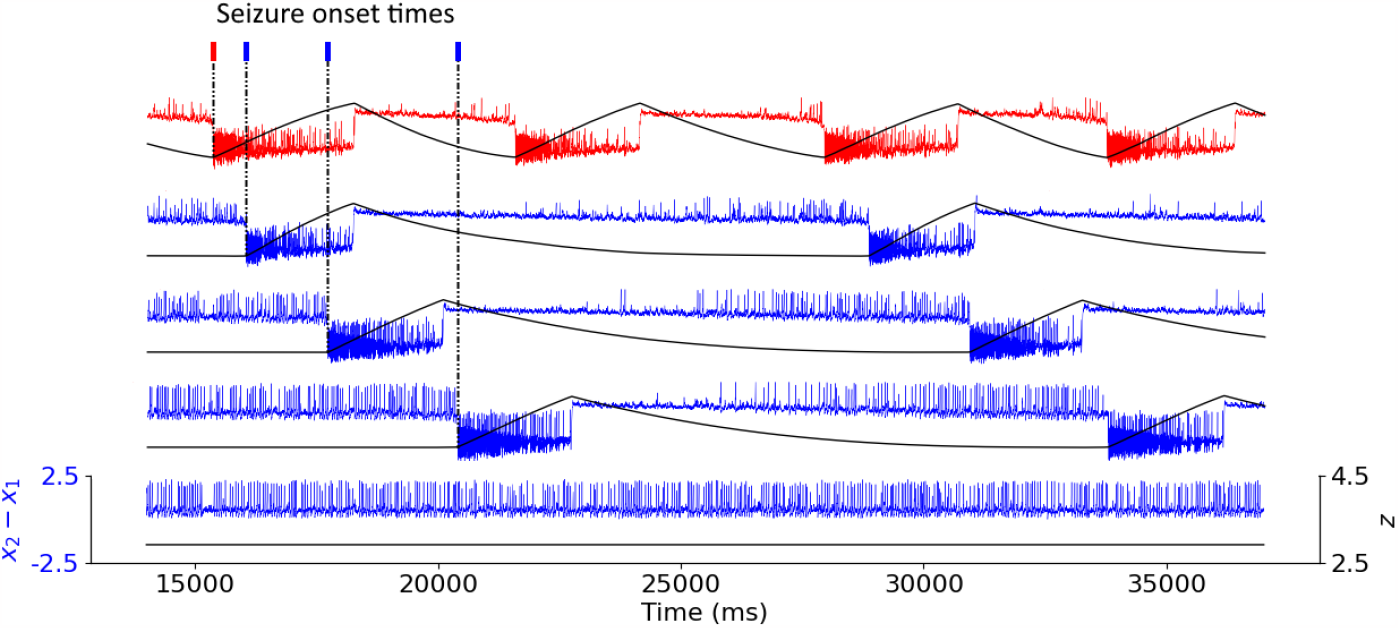
Coupled Epileptor model time-series. Time series of the fast variable *x*_2_*− x*_1_ of the Epileptor model (in color) depicting the neural activity of the nodes, and of the slow variable *z* (in black). The epileptogenic Epileptor (in red) produces spontaneous seizures, with *x*_0_ =*−* 1.6. The non-epileptogenic Epileptors (in blue), with *x*_0_ =*−* 2.1, can either be recruited in the seizure or maintain healthy activity. For all recruited areas, time stamps indicate seizure onset.

### Structural connectome

Our mouse brain network consists of *N* = 98 Epileptors (Melozzi et al., 2017) coupled with a structural connectivity (SC) weight matrix adopted by the Allen Institute that was presented in (Oh et al., 2014) and used in (Melozzi et al., 2017). In the coupling term of Eq. (3), we do not consider self-connections (i.e. *c*_*ii*_ = 0). The SC matrix SC= (*c*_*ji*_) is shown in **Figure** 2, where *c*_*ji*_ denotes the weight of the connection going from area *i* to area *j* in base-ten logarithmic scale for better visualisation. The Allen Institute neuroimaging analysis consistently involves source regions exclusively situated in the right hemisphere. The SC matrix used here the left hemisphere’s counterpart constructed by mirroring the right hemisphere. Originally, the strength of connections between a given region and another was computed as the average across several experiments utilizing those specific brain areas as source and target regions (see (Melozzi et al., 2017) for more details). The SC matrix is divided into four blocks, i.e., R-R, R-L, L-R, and L-L (in a clockwise order from the upper left), symmetries emerge where R-R equals L-L and R-L equals L-R. This assumption is grounded in the notable lateral symmetry observed in the mouse brain, as reported in (Calabrese et al., 2015). The dynamics of Epileptors coupled through this connectome adequately reproduces the seizure recruitment order (in the hippocampus, Subiculum, Enthorinal cortex, Olfactory cortex, Isocortex and Striatum, with epileptogenic left CA1, CA3 and Dentate Gyrus) (Melozzi et al., 2017) that is also experimentally observed by (Toyoda et al., 2013).

**Figure 2.**
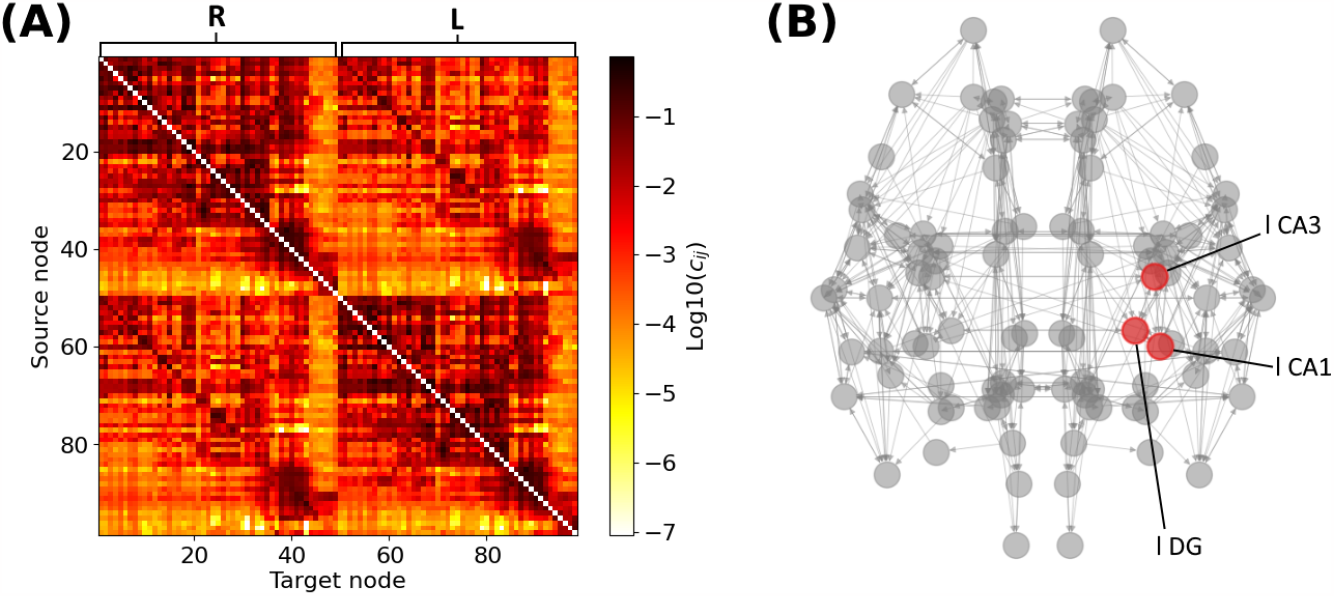
Structural connectivity matrix and mouse brain network. **(A)** Allen Structural Connectivity matrix. The elements of the matrix show the logarithm of the weight for the connections between any pair of brain areas. Node indices ranging from 1 to 49 form the right hemisphere (R), node indices ranging from 50 to 98 form the left one (L). **(B)** Mouse brain network graph. Only connections of weight higher than 0.1 are shown blue for visibility reasons. The red nodes indicate the areas composing the left hippocampus. Note that the left brain hemisphere appears on the right hand side of our template.

For statistical relevance of our computational findings, we also produced a set of 20 connectomes derived from the original SC. Namely, each new weight 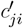 value of the connection between areas *i* and *j* is randomly generated from a normal distribution with mean *c*_*ji*_ and standard deviation 0.1*c*_*ji*_. In the case where a new weight is negative, we set 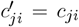. Note that the most prominent modifications occur in the relatively strong connections and that these modifications induced a minor loss of the original SC’s symmetry between left and right hemisphere.

### Network graph measurements

We describe the mice brain network as a graph *G* = (*V, E*) containing a set of vertices *V*, and a set of *E* edges. The adjacency matrix SC= (*c*_*ji*_) is such that *c*_*ji*_ denotes the weight of the connection going from node *i* to node *j*. We characterize each node of the network with different connectivity measurements (see e.g. (Newman, 2010)):

- **Degree**. The amount of connections leaving (resp. arriving) a node *i* is given by its outdegree deg^+^(*i*) (resp. indegree deg^−^(*i*)):

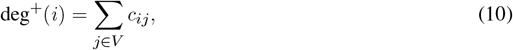

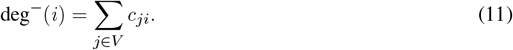

For an unweighted graph, indegree and outdegree represent a number of connections. However, in the case of a weighted graph, they take the real value of the total connective strength the node receives or releases.
- **Eigenvector centrality**. The eigenvector centrality *x*_*i*_ is used to quantify the relative importance of a given node *i*:

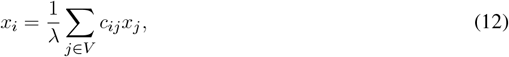

where *λ* is the adjacency matrix’s larger eigenvalue. Eigenvector centrality gets higher when a node has more, stronger connections, especially connections with other central nodes of the network.
- **Average shortest path length**. The shortest path length *l*_*ij*_ between any two nodes *i* and *j* is the total connective length on the shortest path going from one to another, computed following Dijkstra’s algorithm. The length of each connection is artificially defined as *l*_*ji*_ = *c*_*m*_ *− c*_*ji*_, with *c*_*m*_ the largest weight value of a connection in the network, so that strong connections account for short distance. We define the shortest path length of a node *i* as the average shortest path between *i* and any other node of the network:

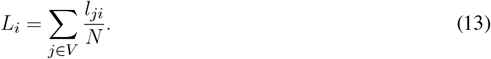

Note that *l*_*ji*_ is always defined as the mice brain graph is complete, i.e., each pair of graph vertices in the graph is connected by an edge. Therefore there is always a way connecting node *i* to node *j*. The lower the shortest path length, the fastest the information goes from one node to another.

To study the relative importance of various nodes within a brain network, these connectivity measurements are normalized. With *m*_*i*_ is the initial connectivity measurement value for node *i* and *m*^max^ the maximal value of the measurement among the brain nodes, we define the normalized value as 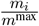 .

## 3 RESULTS

### 3.1 Simulating widespread and localized epileptic seizures

The location of the Epileptogenic Zone in a given mice brain network determines the propagation (i.e. generalized seizure - occurring in both hemispheres of the brain) or non-propagation (i.e., focal - occurring in one area of the brain or hemisphere in the brain that remains localized) of the epileptic seizure. We start by computationally initiating epileptic seizures within different areas of the hippocampus, namely the CA1, CA3 and DG separately and not all three simultaneously (nor combinations of pairs of them). Our rationale in doing so is to begin by investigating whether there is an systematic association between the seizure onset at a certain sub-region (CA1, CA3 and DG) and the type of seizure, e.g. widespread vs localized. **Figure** 3 displays the time distance from the initiation time of a seizure at a EZ node to reach different brain areas in a template of a mice brain slice. **Figure** 3(A) shows an example of a widespread seizure, starting in left-field CA1 and spreading to almost all brain areas (nodes). **Figure** 3(B) shows an example of a focal-localized seizure, starting in left-field CA3. Note that in the latter case, the epileptiform activity remains confined in the vicinity of the EZ. This preliminary finding provides a first hint that an epileptic seizure occurring in CA1 or CA3 may have a rather different propagation in the other brain areas.

**Figure 3.**
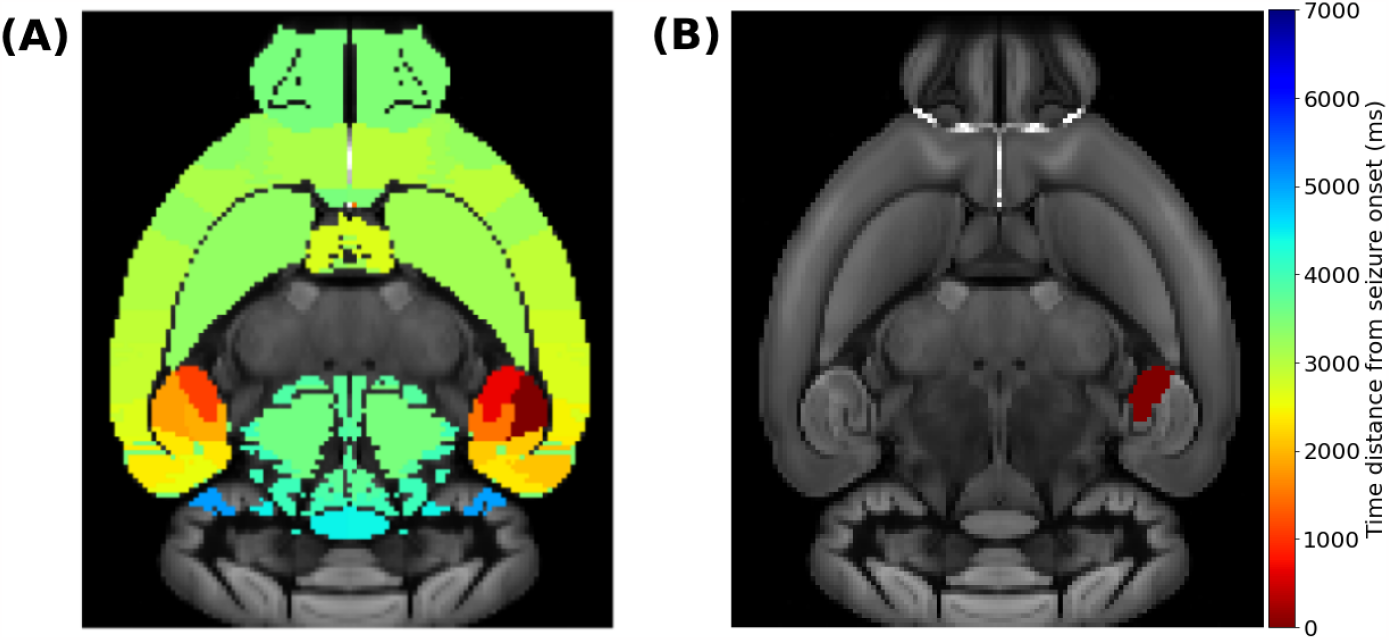
Simulating widespread and localized epileptic seizures on the mouse brain. **(A)** A widespread (generalized) seizure that starts in left-field CA1. **(B)** A local seizure that starts in left-field CA3. The colorbar indicates the time-distance from seizure initiation in each respective brain area during its propagation.

Next, we performed similar simulations by initiating seizures at all available brain areas of the Allen atlas. Our motivation, at this stage, was mainly computational, i.e., to identify other brain areas where the onset of a seizure will result to either a widespread or a localized one. Moreover, and for statistical significance, we repeated the simulation for 20 additional different (but similar) SC matrices that we generated as explained in the Materials and Methods section. These SC matrices are can be found in the Supplementary Material (**Figure** S1 and S2). Note that, as the original connectome has symmetrical connections between left and right hemispheres, the simulations were only performed for EZ within the left hemisphere and were duplicated for the right hemisphere EZ. Regarding the randomly modified SC connectomes, we ran simulations considering EZ regions in both hemispheres separately as the matrix symmetry between the two hemispheres is lost.

**Figure** 4 shows the percentage of seizure-recruited brain areas for each EZ brain area in the original mice brain connectome, labelled as Allen SC in the first row of each panel, and in the 20 SC randomly generated, labelled by their cardinal number (20 rows above the original Allen SC row separated by the while horizontal line). The resulting epileptic seizures allow a clear binary distinction between widespread and local seizures. Namely, our model and the available SC mice connectomes resulted in generating either localized seizures (no widespread propagation at all or limited up to only two regions) or widespread seizures (reaching almost all brain areas). **Figures** 4(A) and (B) summarize the simulations for left and right brain hemispheres respectively. Mild modification in the SC matrices can lead to different types of seizure propagation that are initiated at different EZ areas. Some nodes generate exclusively localized seizures (e.g. areas 1 or 45 in dark blue of the right hemisphere) whereas others generate widespread ones (e.g. areas 7 or 13 in dark red of the right hemisphere). However, there are also EZ regions where the type of the seizure depends on the particular SC matrix (e.g. area 75 in the left hemisphere). The hippocampus area is considered to be among the most common source of epileptic seizures. The yellow frame indicates the three nodes comprising the left hippocampus, namely left-field CA1 (l CA1, node 73), left-field CA3 (l CA3, node 74) and left Dentate Gyrus (l DG, node 75). For all randomized connectomes, left-field CA1 produces widespread seizures, while left-field CA3 produces localized seizures with no propagation. left Dentate Gyrus can either produce local or widespread seizures, depending on the SC matrix.

**Figure 4.**
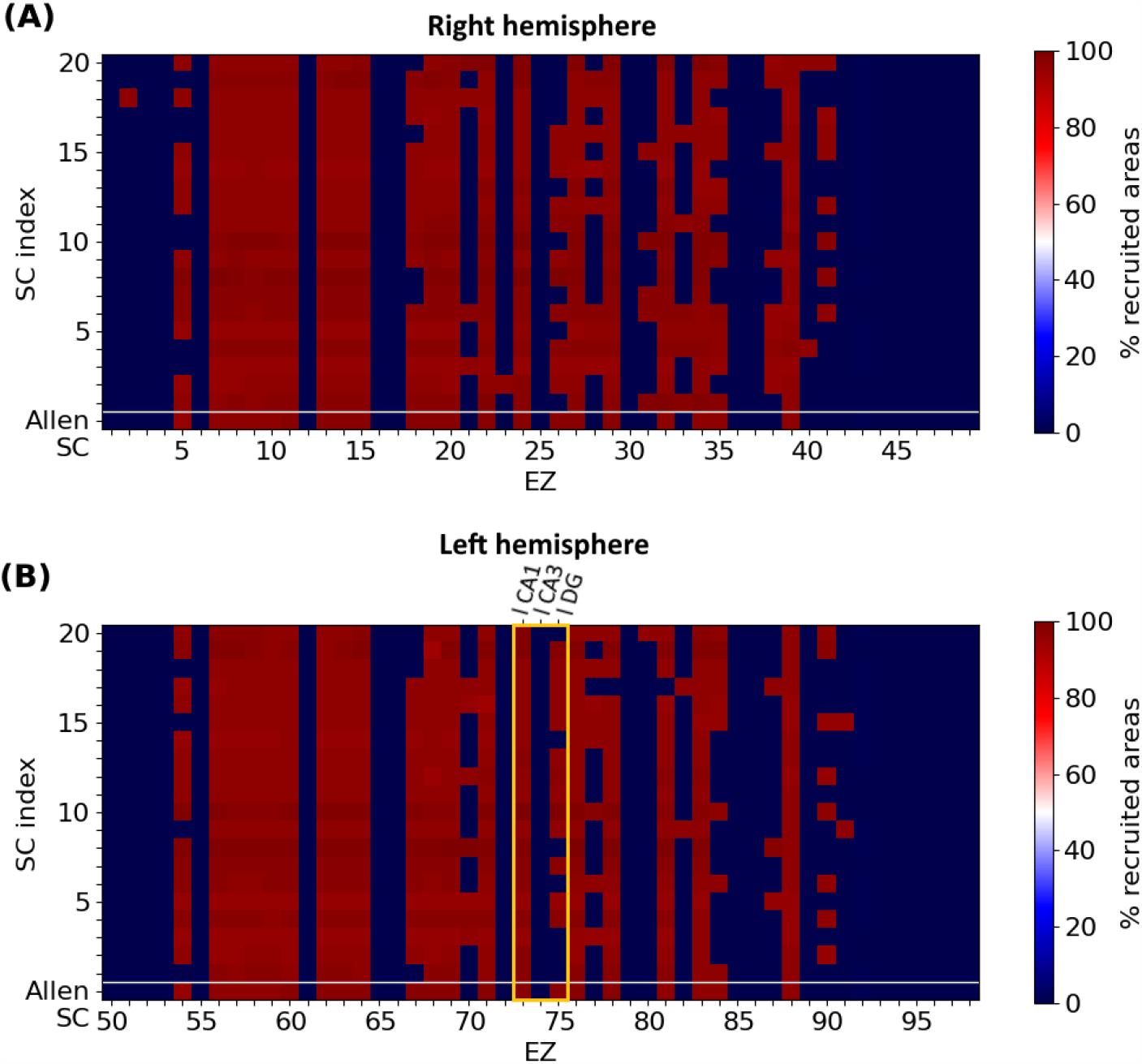
Localized and widespread epileptic seizure for different structural connectomes. We simulated epileptic seizures for the original mice brain connectome (Allen SC, first row of each panel) and for different randomly generated SC matrices (separated by the while horizontal line). **(A)** Right hemisphere and **(B)** left hemisphere. Certain nodes generate exclusively localized or widespread while for some EZ areas the type of the seizure depends on the SC matrix. The yellow frame in panel **(B)** highlights the three nodes comprising the left hippocampal network. Note that our simulations generated either localized seizures (no widespread propagation at all or limited up to only two regions) or widespread seizures (reaching almost all brain areas).

Next, we sought to explore plausible associations between a chosen EZ region’s influence in the network (quantified by a graph connectivity measurement) and the resulted type of seizure (localized vs widespread). **Figure** 5 shows different types of seizure propagation using different SC matrices (initiated at different EZ regions of the Allen atlas). Each panel depicts different combinations of their respective graph connectivity measurements, namely their normalized eigenvector centrality, out-degree, average shortest path length, and the strongest outgoing connection weights. Each point in these plots displays the connectivity properties of a single EZ, and the percentage of brain areas affected during the seizure propagation initiated at this particular EZ. By means of these graph connectivity measurements, we were able to identify three distinct regions delineated by the following thresholds: all EZ whose strongest outgoing connection has a weight value larger than *w*_upper_ = 0.31 produced widespread seizures, while all EZ whose strongest outgoing connection has a weight value lower than *w*_lower_ = 0.22 only produced localized seizures. EZ nodes with values in-between these two values resulted to either localized or widespread seizures (see **Figures** 5 (A)-(D)).

**Figure 5.**
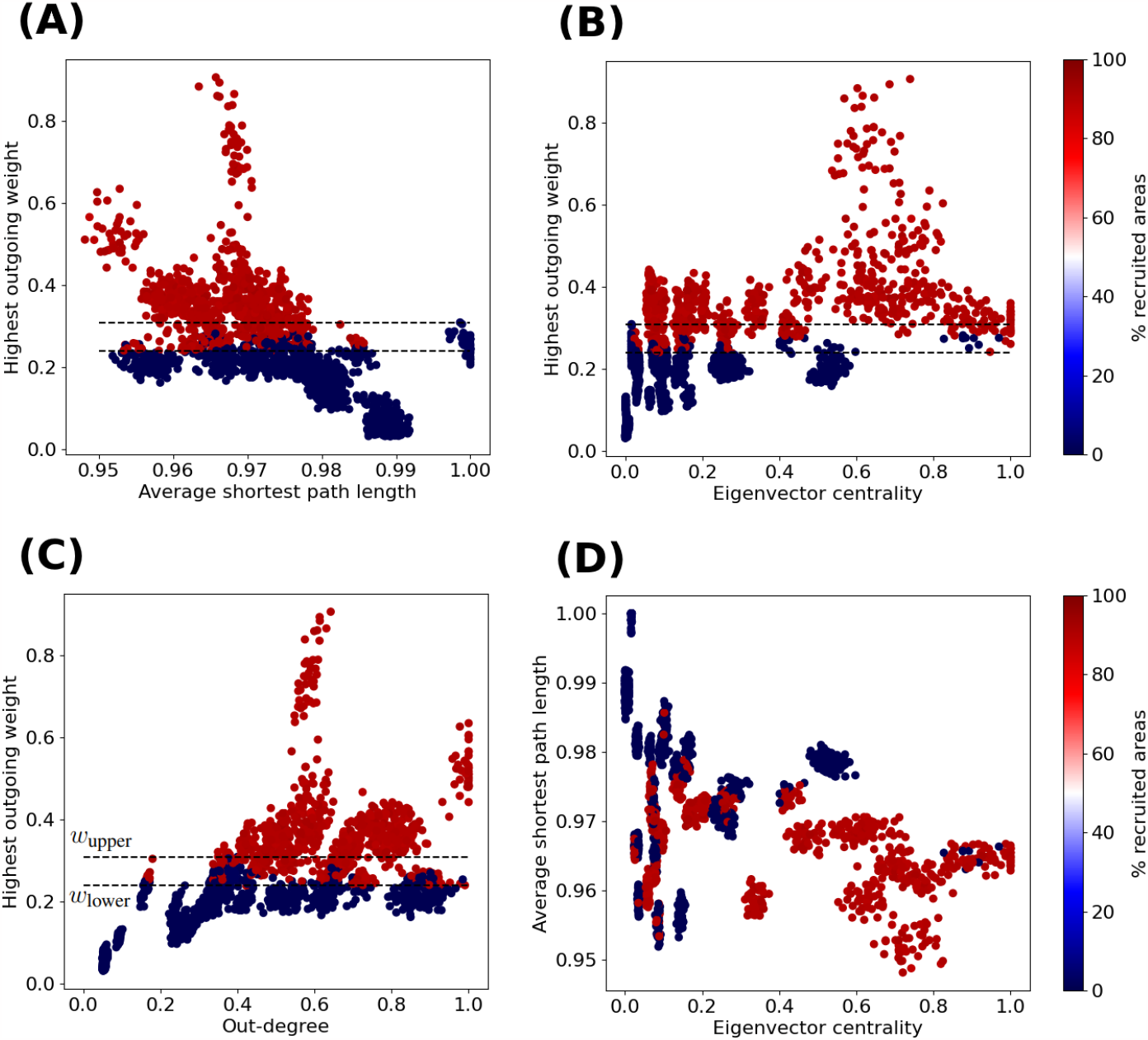
Widespread vs localized epileptic seizures and EZ graph network connectivity measurements. We calculated the normalized average shortest path length, eigenvector centrality and out-degree for each node of the original Allen SC and the 20 additional randomized connectomes (98 nodes per SC matrix). The colormap indicates the fraction of brain areas of a seizure propagation when it is initiated at the respective EZ region (i.e., the points depicted in the plots). Each panel depicts different combinations of their respective graph connectivity measurements, namely the normalized average shortest path length vs the strongest outgoing connection weights **(A)** the eigenvector centrality vs the strongest outgoing connection weight **(B)**, the out-degree vs he strongest outgoing connection weight **C)** and the normalized eigenvector centrality vs the average shortest path length **(D)**. The upper (resp. lower) horizontal dashed line shows the aforementioned thresholds *w*_upper_ (resp. *w*_lower_) of the larger outgoing edge weight (see text for more details).

The eigenvector centrality, out-degree and average shortest path length might also be relevant properties to infer on the node’s (in-)ability to produce a widespread seizure. Clusters of nodes with relatively low average shortest path length value and high eigenvector centrality value (*L*_*i*_ *<* 0.97, *x*_*i*_ *>* 0.30) can systematically produce widespread seizures. On the other hand EZ areas with relatively high average shortest path length value and low eigenvector centrality value result to localized seizures (see cluster near the area with *L*_*i*_ *>* 0.98, *x*_*i*_ *<* 0.20 in **Figure** 5(D)).

### 3.2 Prevention of widespread seizure in Temporal Lobe Epilepsy

#### Connectivity of the hippocampal subnetwork

The hippocampus is a well-known EZ in Temporal Lobe Epilepsy (Toyoda et al., 2013; Buckmaster et al., 2022). In this study, we investigate seizure propagation and confinement when seizure initiation occurs in the different areas of the hippocampus. In the Allen mice brain atlas, the hippocampal subnetwork is composed of three nodes, namely the CA1, CA3 and Dentate Gyrus areas (see **Figure** 2 and the discussion earlier of **Figure** 4). In **Figure** 6(A), we show the hippocampal subgraph. The color of each connection indicates its weight. The hippocampal subnetwork exhibits strong connections within left (resp. right) hippocampus itself, and between left and right hippocampus. The weights of the strong outgoing connection of the left-field CA1 (*w*_l CA1_ = 0.36), left-field CA3 (*w*_l CA3_ = 0.18) and left Dentate Gyrus (*w*_l DG_ = 0.25) strongest outgoing connection are such that *w*_l CA1_ *> w*_upper_ *> w*_l DG_ *> w*_lower_ *> w*_l CA3_. More precisely, the left-field CA1 lies within the connectivity area of widespread seizure production as presented in **Figure** 5. Left DG also lies within the region of potential widespread seizure production, while left-field CA3 is in the region of local seizure production. Note that moreover, CA3 has no strong outgoing connection (with weight higher than 0.1) outside the hippocampus, contrary to left-field CA1 and left Dentate Gyrus. In particular, both left-field CA1 and left Dentate Gyrus present edges leading to left Entorhinal Cortex, lateral part (l ENTl) and left Subiculum (l SUB).

**Figure 6.**
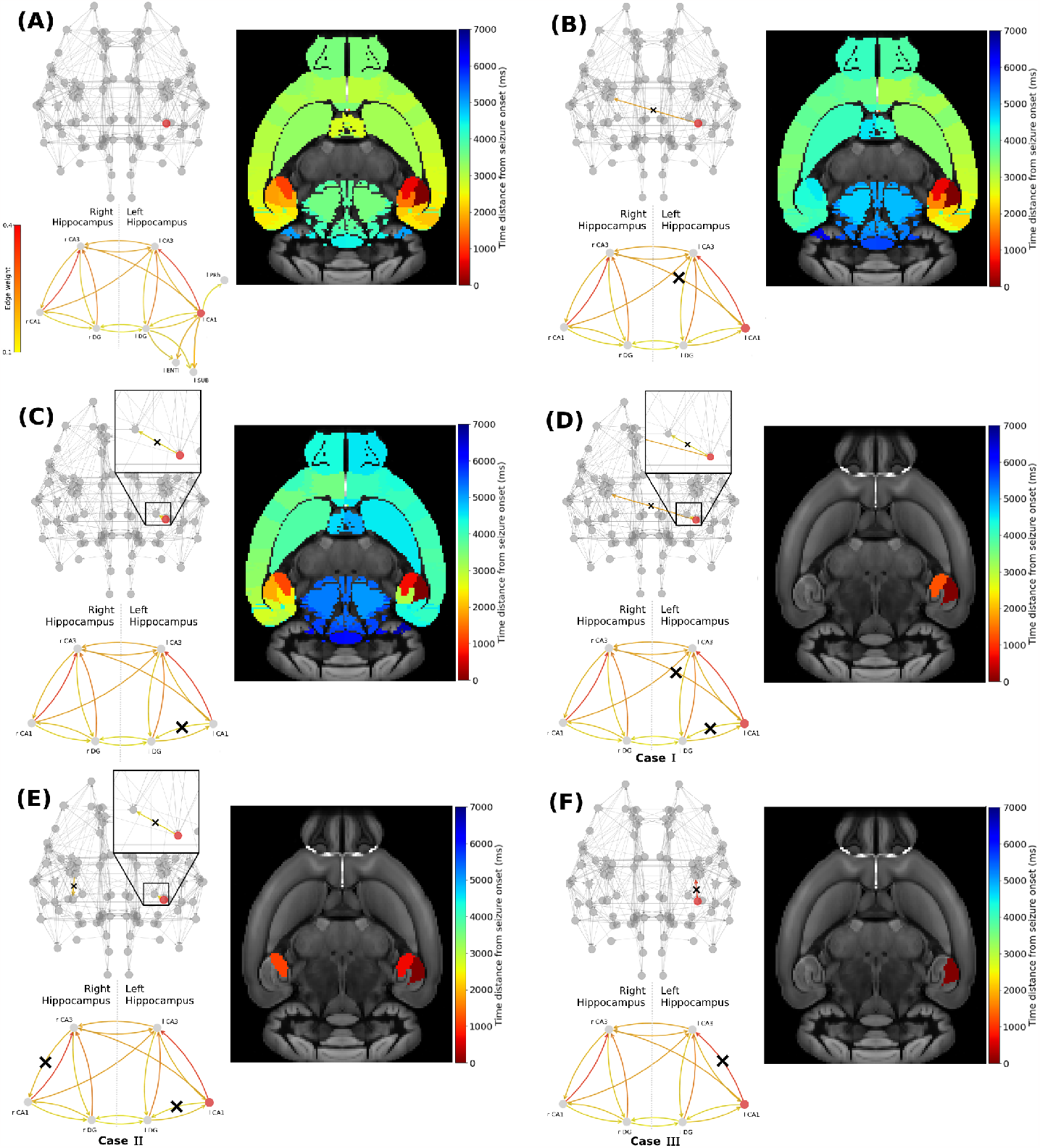
Widespread seizure prevention by edge removal in the left-field CA1.. The upper left panels show the whole brain network the hippocampal subnetwork. Crosses indicate the removed connections, and the edges’ color shows their respective weight. The lower left panels illustrate the left extra-hippocampal connections and their weights values are given in the left colorbar. The right panels show the time distance between seizure initiation in the EZ (left-field CA1) and its onset in each brain area. Panel **(A)** shows an example of a widespread focal seizure propagation in the original Allen connectome, then after removing the inter-hemisphere strong connection **(B)** l CA1 ↛ r CA3, a strong connection within the left hippocampus **(C)** l CA1 ↛ l DG, the inter-hemisphere strong connection and within the left hippocampus **(D)** l CA1 ↛ r CA3 and l CA1 ↛ l DG, allowing inter-hemisphere communication but blocking a strong within the left and right hippocampus **(E)** l CA1 ↛ l DG and r CA3 ↛ r CA1 and blocking the strongest connection and within the left hippocampus **(F)** l CA1 ↛ l CA3.

#### Structural connectome interventions

In order to control wide spread brain seizure propagation, we experimented with two different interventions on the structural connectome to computationally simulate the effects of two clinical approaches for epilepsy. First, we applied *graph edge removal* from relevant EZ nodes to (loosely) simulate a resection-like surgery intervention. Our ultimate goal here is to suppress the communication pathways between certain brain areas which are involved in a widespread seizure by blocking the minimum amount of edges potentially relevant to the propagation. When we remove a given connection from node *i* to node *j*, we set its new weight value in the adjacency matrix to zero and we re-normalize all weights’ values to one (Melozzi et al., 2017; Jehi, 2018; Rubio et al., 2019; Alexandratou et al., 2021). We use the notation *i* ↛ *j*.

In addition, we also focused on an EZ node and we perform a *global outgoing weight reduction* (by some percentage of the initial weight). This (rather simple) approach aims at loosely modeling a suppression of the hyperexcitability of the diseased brain area using for example neuromodulation techniques (Heck et al., 2014; Tsuboyama et al., 2020; Toffa et al., 2020) or the local effect a drug administered in the vicinity of the EZ area (Kanner and Bicchi, 2022). To this end, we decreased the weight values of all outgoing connections from the EZ. Hence, the new weights 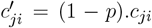, *p* being the percentage of reduction applied. After each of such weight modification, the structural connectivity is re-scaled to preserve its initial total connective strength. We restricted our analysis to the sole EZs left-field CA1 and left Dentate Gyrus regions as the left-field CA3 does not produces widespread seizures (see **Figure** 4(B)).

#### Seizure widespread prevention by edge removal

We use the latter analysis to computationally test the effect of different edge resection options in the original mice brain network on seizure propagation.

### Left-field CA1 edges removal

**Figure** 6 shows the impact of several connections’ removal on the propagation (widespread vs localized) of focal seizures starting in left-field CA1 area. Each upper left panel depicts the modified brain network and the hippocampal subnetwork, with crosses (when present) indicating the removed connections between two or more nodes. Note that for better visualization, we only show connections of weight higher than 0.1. The color of the connections indicates their respective weight values (see also the SC matrix in **Figure** 2). Each lower left panel in **Figure** 6(A), illustrates the hippocampus subnetwork and more particularly the left extra-hippocampal connections and their weights values are given in the left colorbar. Each right panel shows the time distance from seizure initiation in each node of the brain network on the Allen mice brain template. The seizure onset takes place at the dark red region while the end of its propagation at the regions in dark blue. **Figure** 6(A) shows the reference focal seizure initiated in the left hippocampal region and its propagation in the original Allen mice brain connectome. After initiating in left-field CA1, the seizure reaches left-field CA3, then right-field CA3, and propagates rapidly (almost simultaneously) in both hemispheres. The seizure spreads in the left and right hemispheres due to the strong inter-hemispheric hippocampal connections. In **Figure** 6(B), we remove the strongest connection leading from the EZ to the right hemisphere, that is l CA1 ↛ r CA3. Hence, we prevent direct seizure propagation from left to right hippocampus. However, even by doing so, the seizure still spreads eventually into the entire brain network via the left hemisphere following a different evolutionary path and brain areas recruitment sequence. Next, we proceed by following an alternative approach, i.e. we remove a strong connection leading from the EZ to another area of the left hippocampus, namely left Dentate Gyrus. This approach results to a seizure which resembles the activity observed in **Figure** 6(A), namely it reaches both hemispheres. However the propagation turns out to be relatively delayed, longer localized within the left hemisphere before spreading to the right one. In **Figure** 6(D), we present a resection strategy (*Case I*) where we block both inter-hemispheric and intra-hemispheric communication pathways, namely l CA1 ↛ r CA3 and l CA1 ↛ l DG. This approach results to a localized seizure in the vicinity of the EZ area. In **Figure** 6(E), we remove the left intra-hippocampal pathway l CA1 ↛ l DG and search for removal options within the right hippocampus instead of removing inter-hemispheric edges. We find one candidate connection that prevents widespread seizure propagation when being removed, namely the r CA3 ↛ r CA1 (*Case II*). In **Figure** 6(F), we show that the resection of the strongest connection l CA1 ↛ l CA3 (*Case III*) is sufficient to prevent the seizure from spreading to any other area. Note that this is the single EZ outgoing connection with weight higher than the threshold *w*_upper_ identified in **Figure** 5.

### Left Dentate Gyrus edges removal

Next, we conduct a similar analysis with epileptogenic left Dentate Gyrus and show the results in **Figure** 7. After initiating a seizure in left Dentate Gyrus, it spreads to the left hemisphere via left-field CA3 then left-field CA1, and to the right hemisphere via right-field CA3 (see **Figure**7(A)). In **Figure** 7(B), we show that the removal of a strong pathway between the EZ and the right hemisphere hippocampus l DG ↛ r DG has a weak impact on the seizure widespread propagation. In **Figure** 7(C), we show that the removal of a strong connection leading from the EZ to other within the left-hippocampal areas l DG ↛ l CA1 results in a delayed widespread seizure propagation indicated by the blue color on the template. Then, we present three examples that yield to seizure localization. Namely, in **Figure** 7(D) we removed strong connections between the EZ and another left-hippocampus area, and between left and right hippocampus. l DG ↛ l CA1 and l DG ↛ r DG (*Case I’*). In **Figure** 7(E) we removed strong connections leading from the EZ to other parts of the left hippocampus and from right-field CA3 to other parts of the right hippocampus l DG ↛ l CA1 and either r CA3 ↛ r DG or r CA3 ↛ r CA1 (*Case II’*). And finally in **Figure** 7(F) we removed the strongest connection l DG ↛ l CA3 (*Case III’*). We have also experimented with other similar effective strategies for focal epilepsy localization that can be found in the Supplementary Material (**Figure S3**).

**Figure 7.**
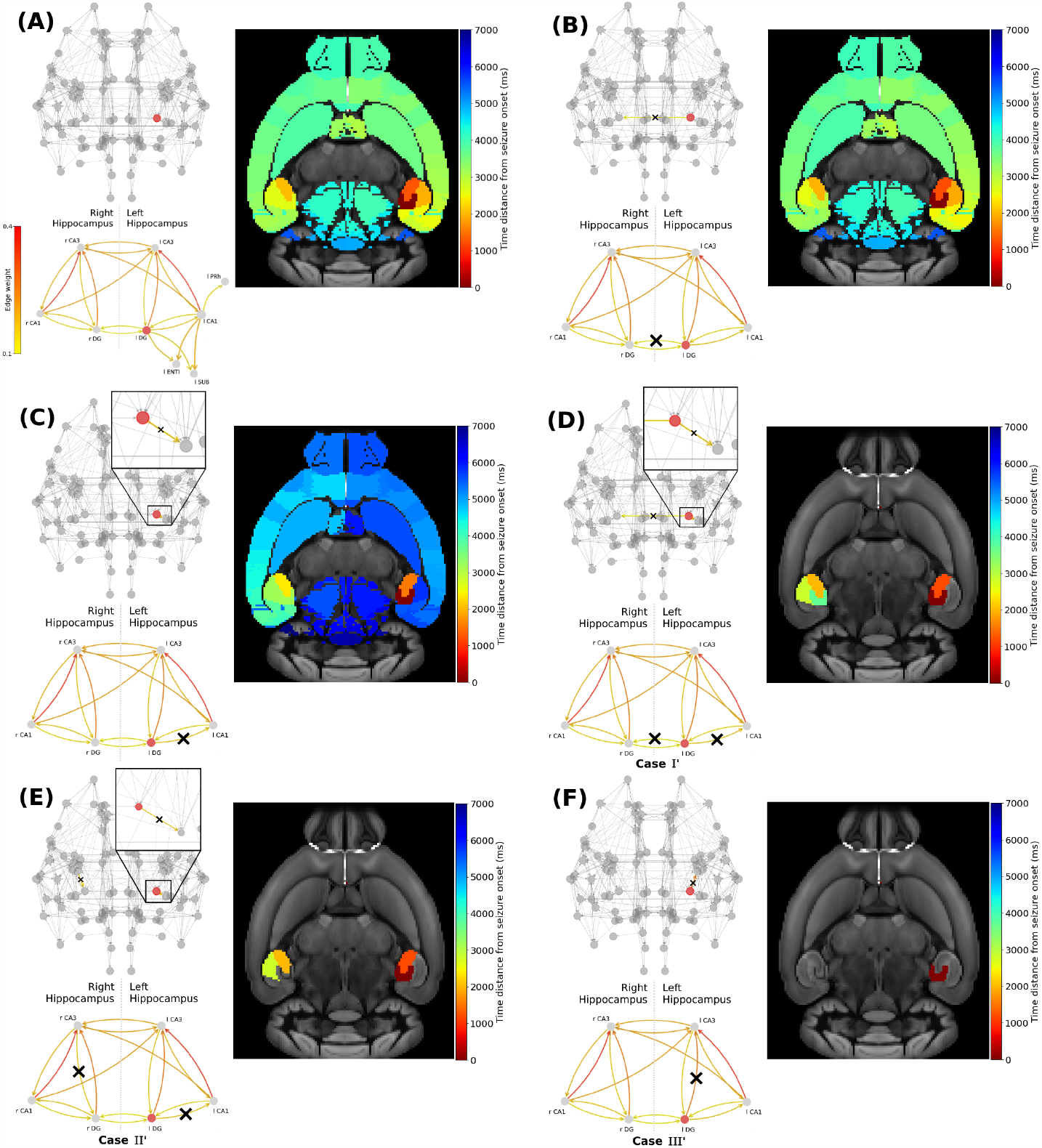
Widespread seizure prevention by edge removal in the left DG. The upper left panels show the whole brain network the hippocampal subnetwork. Crosses indicate the removed connections, and the edges’ color shows their respective weight. The lower left panels illustrate the left extra-hippocampal connections and their weights values are given in the left colorbar. The right panels show the time distance between seizure initiation in the EZ (left-field CA1) and its onset in each brain area. Panel **(A)** shows an seizure propagation in the original Allen connectome. Following a similar approach in removing graph edges (inter-hemisphere and/or within the left/right hippocampus) we gradually proceed by removing **(B)** l DG ↛ r DG, **(C)** l DG ↛ l CA1, **(D)** l DG ↛ r DG and l DG ↛ l CA1, **(E)** l DG ↛ l CA1 and r CA3 ↛ l DG and **(F)** l DG ↛ l CA3.

In **Figures** 6 and 7, we illustrated the result of the aforementioned edge removal approaches using the original Allen mouse brain connectome. In order to further validate our findings, we also ran simulations with the additional SC matrices generated by the original Allen SC matrix as describe on the Materials and Methods section. By doing this, we aimed to improve the statistical significance of our results. **Figure** 8 shows the percentage of recruited areas when applying the latter resection options to the different connectomes, where the left-field CA1 (**Figure** 8(A)) and left DG (**Figure** 8(B)) is the EZ respectively. In both cases, the single resection of EZ ↛ l CA3 (*Case III* and *III’*) is sufficient to achieve seizure confinement in the EZ in all connectomes. Starting with the left-field CA1 as an EZ, in **Figure** 8(A), we show that removal edge *Case I* (second column, l CA1 ↛ r CA3 and l CA1 ↛ l DG, see **Figure** 6(D)) and *Case III* (forth column, l CA1 ↛ r CA3 and l CA1 ↛ l DG, see **Figure** 6(F)) approaches systematically prevent seizure widespread propagation for all SC matrices. Removal edge *Case II* (third column, l CA1 ↛ l DG and r CA3 ↛ r CA1, see **Figure** 6(E)) effectively prevents widespread seizure propagation for 18 out of the 20 SC matrices. In **Figure** 8(B), we present a similar analysis when the EZ is now the left Dentate Gyrus area and when implementing the edge removal *Case I’* (**Figure** 6(D)), *Case II’* (**Figure** 6(E)) and *Case III’* (**Figure** 6(F)). The first two resulted in widespread seizure propagation for 18 out of the 20 SC matrices while the latter one to localized seizure for all connectomes.

**Figure 8.**
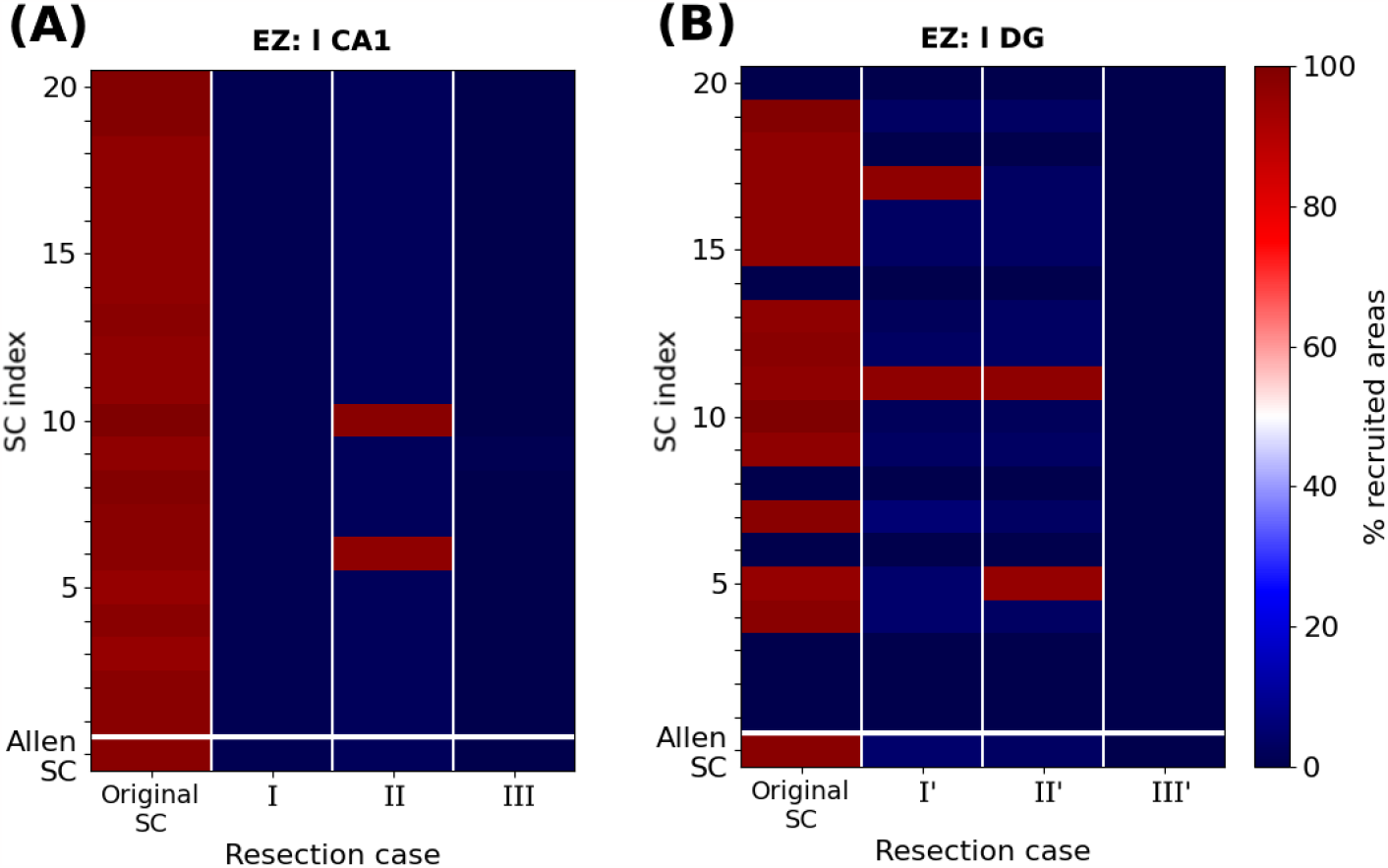
Percentage of recruited areas after edge removal for different mice connectomes. **(A)** Simulations of a focal epileptic seizure initiated in the left-field CA1 area for the original SC Allen weight matrix (first row) and 20 additional SC ones (see text for more details), The first column (dark red) shows the robustness of the finding presented in **Figure** 6 (A), i.e. that the left-field CA1 area generates systematically widespread seizures. Removal edge *Case I* and *III* approaches both systematically yield seizure localization (dark blue) while in *Case II* two connectomes (red color) yield to widespread seizure propagation and the rest to localized (blue color). **(B)** Simulations of a focal epileptic seizure initiated in the left DG area for the original SC Allen weight matrix and 20 additional SC ones and the edge removal *Case I’, II’* and *III’* approaches (see **Figure** 7).

#### Seizure widespread prevention by edge outgoing weight reduction

We introduce an alternative procedure to prevent widespread seizure propagation. Instead of removing network connections, we mimic the reduction of EZ hyperexitability with a global decrease of its outgoing weight, the extreme case of which would be the suppression of any EZ output. In terms of connectivity measurements, outgoing weight reduction has a direct impact on the EZ’s out-degree and the weight of its strongest outgoing edge, and by extension on its eigenvector centrality. To identify a minimal level of reduction one should apply to prevent seizure propagation, we perform simulations of an epileptic seizure starting in left-field CA1, then in left Dentate Gyrus, before any reduction and after the reduction of 10%, 20%, 30%, 40% and 50% of the EZ’s outgoing edges weights.

**Figure** 9 shows the percentage of recruited areas when applying the latter reductions on the different randomized connectomes. For a seizure starting in left-field CA1, the control gets efficient in most of the cases when reducing the EZ’s outgoing weights of 30%. We achieve seizure confinement in all connectomes by reducing the EZ’s outgoing weights of at least 40%. For a seizure starting in left Dentate Gyrus, confinement is achieved in all connectomes with a reduction of 10%.

**Figure 9.**
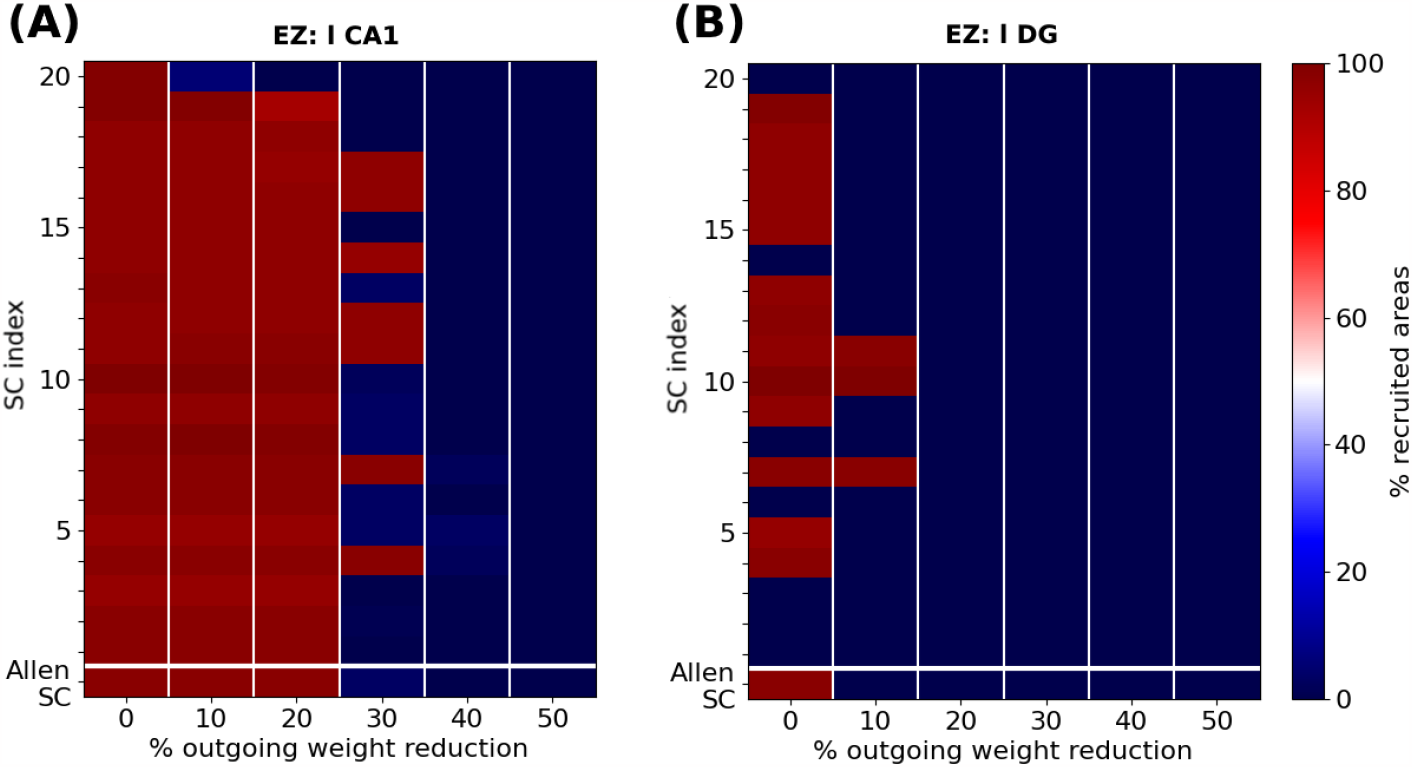
Percentage of recruited areas for different levels of outgoing weight reduction. Simulations of a focal epileptic seizure initiated in **(A)** the left-field CA1 area for the original SC Allen weight matrix (first row) and 20 additional SC ones (see text for more details). Outgoing weight reduction of at least 40% systematically yields seizure localization (dark blue). **(B)** Simulations of a focal epileptic seizure initiated in the left DG area for the original SC Allen weight matrix and 20 additional SC ones. Outgoing weight reduction of at least 20% systematically yields seizure localization.

Finally, we show the effect of the latter resection and outgoing weight reduction strategies on the EZ’s connectivity.

**Figure** 10 shows the eigenvector centrality and larger outgoing weight value of **(A)** left-field CA1 and **(B)** left Dentate Gyrus in the Allen connectome, before and after performing structural modifications. The horizontal dashed lines indicate the upper and lower threshold values for the larger outgoing weight *ω*_*upper*_ and *ω*_*lower*_ defining regions of widespread and local seizure production (see **Figure** 5). The color of the nodes indicate the percentage of seizure-recruited areas in each scenario. Along the long diagonal arrow, we show from top to bottom the effect of the EZ outgoing weight reduction of: 0% (Original SC), 10%, 20%, 30%, 40% and 50%.

**Figure 10.**
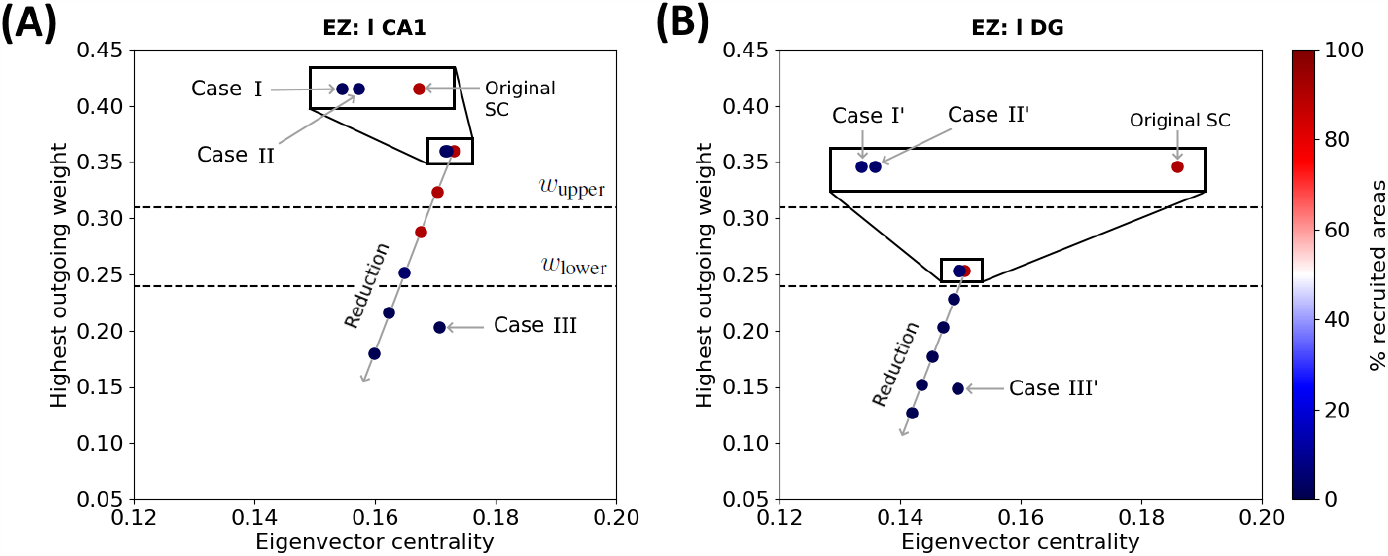
EZ connectivity and percentage of seizure-recruited areas after outgoing weight reduction and edge resection. Each point shows the eigenvector centrality and the largest outgoing weight value in the Allen connectivity of **(A)** left-field CA1 and **(B)** left Dentate Gyrus, before and after applying outgoing weight reduction and edge removal strategies *Case I, II* and *III* (resp. *Case I’, II’* and *III’*) as labelled in **Figure** 6 and **Figure** 7. The points aligned on the diagonal arrow represent, from top to bottom, an outgoing weight reduction of 0% (Original SC), 10%, 20%, 30%, 40% and 50%. The color of each point shows the percentage of recruited brain areas when the seizure starts in the corresponding node. The two dashed lines show the upper (resp. lower) threshold of the largest outgoing weight from separating the widespread from localized seizures.

In **Figure** 10(A), we present the effects of edge removal approaches *Case I* (l CA1 ↛ r CA3 and l CA1 ↛ l DG), *Case II* (l CA1 ↛ l DG and r CA3 ↛ r CA1) and *Case III* (l CA1 ↛ l CA3) for the left-field CA1 area and only for the original Allen SC Matrix. When applying outgoing weight reduction, the progressive decrease of the EZ’s outgoing connective strength leads its connectivity towards lower eigenvector centrality values, and especially lower strongest outgoing weight values. The approach *Case III* also results in reducing the EZ’s strongest outgoing weight to value below *ω*_lower_ threshold. Note that EZ nodes with outgoing weight values lower than *ω*_lower_ yield to seizure localization. The edge removal approaches *Case I* and *II* also mildly reduce the EZ’s eigenvector centrality, and lead to seizure localization without any modification of its strongest outgoing connection. In **Figure** 10(B), we present the effects of the edge removal approaches *Case I’* (l DG ↛ r DG and l DG ↛ l CA1), *Case II’* (l DG ↛ l CA1 and r CA3 ↛ r DG) and *Case III’* (l DG ↛ l CA3) for the left Dentate Gyrus and same connectome. The overall trend is again similar, namely focal seizures remain localized as the outgoing weight value of the node decreases after our intervention and crosses the lower threshold (lower dashed line). Note that *Case III’* has a strong impact in reducing the strongest outgoing connection weight in left Dentate Gyrus while resections *Cases I’* and *II’* cause an eigenvector centrality value reduction. A similar Figure with all SC matrices and analysis can be found in the Supplementary Material (**Figure S4)** .

### Brain functionality after edge resection and reduction

Modifying the brain network structure induces changes in functional connectivity, leading to either enhanced or reduced functional couplings between different brain areas. The objective is to minimize connectome interventions to avoid drastic alterations in brain functionality. For each approach involving the removal of edges and weight reduction, we computed the modified simulated Functional Connectivity (FC) matrix using the Epileptor model. This matrix was then compared to the FC matrix generated by the original Allen Structural Connectivity (SC) matrix for the healthy mouse. The corresponding figures illustrating these comparisons can be found in the Supplementary Material (**Figure S5**). Our approaches aim to minimize structural modifications compared to standard (also clinical) procedures in the literature, which involve the removal of the entire left and right hippocampus.

## 4 DISCUSSION

In this study we employed the Epileptor model to simulate widespread and localized epileptic seizures (**Figures** 1 and 3) and investigate the relationship between the location of an EZ, its connectivity graph significance in the network (**Figures** 4 and 5), and the widespread propagation of such seizures in a mouse brain structural connectome (**Figure** 2). When initiating focal seizures within the three hippocampal sub-regions (from the Allen atlas) we observed that the left-field CA1 area systematically generates widespread seizures, the left-field CA3 systematically generates localized seizures, and left Dentate Gyrus can generate both widespread and localized seizures, depending on the SC weight matrix (**Figures** 4).

Our next objective was to identify plausible and effective strategies that are able to prevent widespread seizures. To achieve that, our first approach (**Figures** 6 and 8) was to selectively remove a minimal amount of EZ node edges (targeted disruption of specific network connections). Instead of adopting the approach of computationally “resecting’ the entire EZ tissue from both brain hemispheres and as performed in surgical settings (see e.g., (Jirsa et al., 2016; Melozzi et al., 2017; An et al., 2019; Olmi et al., 2019; Nissen et al., 2021)), here we aimed to systematically identify and eliminate (or block) the minimal set of connections necessary to prevent generalised seizures. In our second approach (**Figures** 7 and 9), we attempted to computationally (and somewhat loosely) model the impact of a drug (Kanner and Bicchi, 2022) or a neuromodulation approach (suppression of network hyperexcitability) in the vicinity of an EZ region (see e.g. (Tsuboyama et al., 2020)). Therefore, we adjusted the outgoing weight connections of an EZ node in our structural connectome to emulate the inhibitory effect of such a drug or external stimulation in the proximity of the targeted area around the EZ. In both approaches, our ultimate goal is to minimize the surgical or medical intervention while preserving as much as possible the pre-surgical original structural connectivity and brain functionality. Our findings indicate that strong inter and intrahippocampal nodes’ communication is important for the widespread seizure propagation. The successful (or not) outcome in preventing widespread seizure propagation can be computationally associated with a threshold value crossing of the larger weight outgoing of the EZ (left-field CA1 or left Dentate Gyrus) area caused by our interventions (in silico) as shown in **Figure** 10 (see also **Figure S4** in the Supplementary Material where we show the results from SC matrices used for each approach).

The research into the structural organization of the brain has enhanced our understanding of the processing and integration of information through specialized neural circuits distributed across the brain. The application of current network theory to brain connectomes has played a crucial role in this advancement, revealing a set of universal organizational principles that govern brain connectivity. These principles seem to be consistent across different species and scales (see e.g., (van den Heuvel and Sporns, 2019). In (Coletta et al., 2020), the authors used high-resolution mapping of the mouse axonal connectome to uncover novel foundational wiring principles in the mammalian brain, providing a detailed understanding of how neural information is processed and transmitted across different spatial scales. They employed a voxel-level description which revealed organizational principles such as the directional separation of hub regions into neural sink and sources. In a next step, these findings could be taken into account for the graph analysis.

Computational studies have emerged as a powerful approach to advancing our understanding of seizure suppression mechanisms (Taylor et al., 2014), offering insights that bridge the gap between theoretical modeling and clinical intervention. A multitude of recent scientific articles have contributed to this burgeoning field. For instance, researchers have explored the potential of robust control strategies for deep brain stimulation in childhood absence epilepsy, as demonstrated in the work by (Rouhani et al., 2023). Utilizing real-world data, (Brogin et al., 2023) presented a computational framework for the identification and control of epileptic seizures, showcasing the applicability of computational methodologies to real-life scenarios. Furthermore, the prediction of seizure suppression effects through computational modeling has been exemplified by (Ahn et al., 2017), underlining the potential of simulation-based approaches in assessing treatment outcomes. Our work is a step forward into the understanding on how to optimize the impact of such stimulation.

Intricate dynamics of seizure termination and postictal EEG suppression have been uncovered through computational investigations, as highlighted by (Bauer et al., 2017), shedding light on the mechanisms governing convulsive seizures. The design of patient-specific neurostimulation patterns for seizure suppression has also been a subject of study, as evidenced by (Sandler et al., 2018), offering tailored solutions to individual patients. Additionally, the interplay between neural activity and ion concentration changes in localized seizures has been elucidated by (Gentiletti et al., 2022), stressing the critical role of computational methods in deciphering complex neurophysiological processes. Nowadays there are already several devices (approved by the U.S. Food and Drug Administration) aiming to reduce the frequency of seizures, namely Vagus Nerve Stimulation (VNS), Responsive Neurostimulation (RNS), and Deep Brain Stimulation (DBS) (see (Skrehot et al., 2023) and references therein for a recent review). Here again the weight decrease may give a hint on the location of the stimulation efficiency.

The clinical significance of epileptic seizure propagation lies primarily in the fact that the epileptic manifestations cannot be solely attributed to the activity within the seizure focus itself; rather, they result from the spread of epileptic activity to other brain structures. Propagation, particularly when leading to secondary generalizations, poses a significant risk to patients, including recurrent falls, traumatic injuries, and unfavorable neurological outcomes. Anti-seizure medications (ASMs) exert diverse effects on propagation with varying potencies. Notably, for individuals resistant to drugs, targeting seizure propagation may enhance quality of life even without a substantial reduction in simple focal events (see e.g. (Brodie et al., 2016; Khateb et al., 2021).

Network hyperexcitability is a potential contributor to cognitive dysfunction in individuals also with Alzheimer’s disease (AD). In AD patients, hyperexcitability and epileptiform activity are frequently observed and have been linked to impaired cognitive function. Studies conducted on transgenic mouse models of AD in preclinical settings have shown that suppressing epileptiform activity with anti-seizure drugs is associated with improved behavior and a reduction in histopathological indicators of chronic network hyperexcitability in the hippocampus. Drugs such as Levetiracetam (LEV) are commonly used as anti-seizure medication and have been reported to effectively suppress epileptiform spikes and enhance synaptic and cognitive function in mouse models of AD. Moreover, LEV is currently undergoing human trials (in both adults and children) for the treatment of seizures and long-term epilepsy, as indicated by research studies such as (Vossel et al., 2021) and (Onos et al., 2022).

The intricate interplay between excitation and inhibition is fundamental to the proper functioning of neural circuits within the brain. The excitation-inhibition balance, characterized by the equilibrium between excitatory and inhibitory inputs onto neurons, plays a crucial role in shaping the structural connectivity between different brain areas (Bergoin et al., 2023). Changes in this balance have been linked to alterations in the strength of synaptic connections affecting the synaptic and structural plasticity, thereby influencing the overall network architecture and information processing capabilities of the brain. For instance, studies have demonstrated that an excessive excitation-inhibition ratio can lead to aberrant plasticity and altered synaptic weights, potentially contributing to conditions such as epilepsy (see e.g., (Huberfeld et al., 2007). Conversely, a decrease in excitation-inhibition balance has been associated with disrupted neural synchrony and impaired cognitive functions, highlighting the delicate nature of this equilibrium (Roopun, 2008; Yizhar et al., 2011).

Recent research has focused extensively on understanding brain activity at a large scale using resting-state functional MRI (rs-fMRI). The conventional approach typically considers structural connectivity (SC) and the functional connectivity (FC) separately. However, this oversimplification ignores the dynamic engagement of white matter tracts during specific tasks. A more refined concept, termed “resting-state informed structural connectivity” (rsSC), has been introduced to incorporate information from rs-fMRI and infer the dynamic white matter engagement specific to the brain’s state. The resulting rsSC, or resting-state informed structural connectome, reveals the structural network underlying observed rs-fMRI correlations. This approach detects alterations in rsSC community structure in diseased subjects compared to controls. Notably, the original setup does not infer the “directionality” of white matter tracts as either “excitatory” or “inhibitory.” The incorporation of the co-activation (excitatory) or silencing (inhibitory) effects into a hybrid rsSC framework that can allow to infer the brain’s E-I balance can be found in (Ajilore et al., 2013; Fortel et al., 2019, 2020, 2022, 2023) (see also (Manos et al., 2023) for a recent study demonstrating the advantages of using rsSC in modeling time series in whole brain dynamics).

The insights accumulated from computational studies pave the way for more targeted and effective approaches to managing epilepsy. From investigating the impact of spiking timing stimulation on frequency-specific oscillations (Quinarez et al., 2023) to exploring the potential of linear delayed feedback control in thalamocortical models (Zhou et al., 2020), these studies collectively contribute to an expanding body of knowledge that spans from the cellular level (Lu et al., 2017) to network dynamics (Depannemaecker et al., 2021). Moreover, the translation of computational findings into clinical practice has been deliberated upon by (Brinkmann et al., 2021), emphasizing the promising trajectory of computational seizure forecasting. As the field continues to grow, collaboration between computational neuroscientists and medical practitioners holds the potential to revolutionize our ability to suppress seizures and improve the quality of life for individuals living with epilepsy.

Moreover, similar simulations are already being used in computational studies as a tool during pre-surgical stages in the identification of epileptic zones for patients who undergo surgery, In (Makhalova et al., 2022; Jirsa et al., 2023), the authors employed a computational brain modeling method, the so-called Virtual Epileptic Patient (VEP), informed by stereoelectroencephalography (SEEG), and anatomical personalized data to simulate seizures in drug-resistant epilepsy patients. The retrospective analysis of 53 patients revealed that VEP demonstrated higher precision in detecting the EZ compared to clinical analysis and the overall prediction of seizure-free outcomes.

Our study has of course some limitations: First, our analysis was based on the Epileptor model and the Allen mouse atlas with a fixed granularity. To partially mitigate this weincorporated the generation of additional SC matrices to account for some variability and to enhance the statistical significance of our results. Moreover, we did not consider delays in our simulations and we kept all the model parameters and the external noise term fixed throughout our study. However, before choosing the epileptogenicity parameter value *x*_0_ for the EZ regions, we carried out several simulations with other values relatively close to the chosen value and we found rather similar global seizure (widespread or localized) propagation effects to those reported in our manuscript. A similar analysis can be extended for a human connectome in the future. That is to investigate whether similar seizure onset conditions lead to similar types of seizure evolution and whether the intervention approaches we tested here are also effective to a human connectome (network). Our work is largely computationally driven and, at this stage at least, may not immediately provide a direct therapeutic protocol for clinical implementations. However, we believe that it can shed some light on the subject and with follow-up studies help in gaining a better understanding of the underlying mechanisms leading to the onset of different types of seizures and their prevention.

## CONFLICT OF INTEREST STATEMENT

The authors declare that the research was conducted in the absence of any commercial or financial relationships that could be construed as a potential conflict of interest.

## AUTHOR CONTRIBUTIONS

JC: Methodology, Software, Formal analysis, Writing - original draft, review and editing, Visualization. MQ: Methodology, Review and editing, Supervision. YT: Methodology, Review and editing, Supervision. TM: Conceptualization, Methodology, Resources, Writing - original draft, review and editing, Supervision, Project administration, Funding acquisition.

## FUNDING

JC was supported by the LABEX MME-DII (ANR-16-IDEX-0008) PhD grant and CY cognition grant of the CY Cergy-Paris University. TM was supported by the Labex MME-DII (ANR-11-LBX-410 0023-01) French national funding program. MQ was supported by a CNRS financing for a research semester in the IPAL Lab in Singapore. YT was supported by UKRI MRC Project Grant MR/T002786/1. We also acknowledge the use of the Osaka compute-cluster resources of CY Cergy-Paris University.

## ACKNOWLEDGMENTS

We would like to thank Viktor Jirsa, Alex Leow, Damian Battaglia, Sandra Diaz-Pier, Liang Zhan and Giovanni Rabuffo for several fruitful discussions. Juliette Courson would like to thank the support by The Virtual Brain framework community for all the help they provided regarding the software setup and fine tuning.

## DATA AVAILABILITY STATEMENT

The original contributions presented in the study are included in the article/supplementary material, further inquiries can be directed to the corresponding author.

## Supplementary Material

### 1 ADDITIONAL STRUCTURAL CONNECTIVITY MATRICES GENERATED FROM THE ALLEN CONNECTOME

In **Figure** S1, we show in a logarithmic scale the SC of the 20 generated SC. We preserve the structural properties of the Allen connectome (see **Figure** 1 in the main article). In **Figure** S2, we display the difference between the original weight *c*_*ji*_ and the new weight 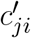 of their connections. Red elements indicate strengthened connections. Blue elements indicate weakened connections. While conserving the overall structure of the SC, there is a significant change in some of the strongest edges weight.

### 2 NETWORK EDGE REMOVAL

In **Figure** S3, we show two resection procedures for the localization of epileptic seizures initiating in l CA1. In **Figure** S3(A), we remove the inter-hemispheric pathway l CA1 ↛ r CA3 and the pathways leading from the EZ to extra-hippocampal left areas, namely left Entorhinal Cortex (l ENTl) and left Subiculum (l SUB). In **Figure** S3(B), we remove the inter-hemispheric pathway l CA1 ↛ r CA3 and the pathways leading from l ENTl, l SUB to high-centrality nodes of the brain network, namely the left Ectorhinal Cortex (l ECT), Perirhinal Cortex (l PRh) and Temporal Association (l TEa) areas. With these two procedures, we achieve seizure localization in a reduced number of brain areas, which suggests that network hubs are important for epileptic seizure propagation.

In **Figure** S4(A), we present the effects of outgoing weight reduction and edge removal approaches *Case I* (l CA1 ↛ r CA3 and l CA1 ↛ l DG), *Case II* (l CA1 ↛ l DG and r CA3 ↛ r CA1) and *Case III* (l CA1 ↛ l CA3) for the left-field CA1 area in the 20 generated randomization of the Allen connectome. In **Figure** S4(B), we present the effects of outgoing weight reduction and edge removal approaches *Case I’* (l DG ↛ r DG and l DG ↛ l CA1), *Case II’* (l DG ↛ l CA1 and r CA3 ↛ r DG) and *Case III’* (l DG ↛ l CA3) for the left Dentate Gyrus area in the 20 generated randomization of the Allen connectome. The two dashed lines show the upper (resp. lower) threshold of the largest outgoing weight separating the widespread from localized seizures (see **Figure** 10 and relevant discussion in the main manuscript for more details). When applying outgoing weight reduction, the progressive decrease of the EZ’s outgoing connective strength leads its connectivity towards lower eigenvector centrality values, and especially lower strongest outgoing weight values. In all 20 generated SC matrices, seizure localization is achieved when the EZ passes below the *ω*_*lower*_ threshold. The approach *Case III* also results in reducing the EZ’s strongest outgoing weight to value below *ω*_lower_ threshold. The edge removal approaches *Case I* and *II* (resp. *Case I’* and *Case II’* mildly also reduce the EZ’s eigenvector centrality, and most of the time leads to seizure localization without any modification of its strongest outgoing connection.

### 3 FUNCTIONAL CONNECTIVITY BEFORE AND AFTER STRUCTURAL INTERVENTIONS

We also use functional analysis to evaluate time correlation between spatially distant brain areas’ activity. To simulate functional connectivity, we compute 10 minute-long time-series of exclusively non-epileptogenic Epileptors coupled through the SC, to reproduce normal brain activity. The first 10 seconds of these time-series are removed to discard the initial bursting due to random initial conditions (the system requires some time to reach its stable state due to transient effects). The Epileptor activity is converted into simulated BOLD activity time-series using hemodynamic response functions implemented in TVB (Friston et al., 2000).

**Figure S1.**
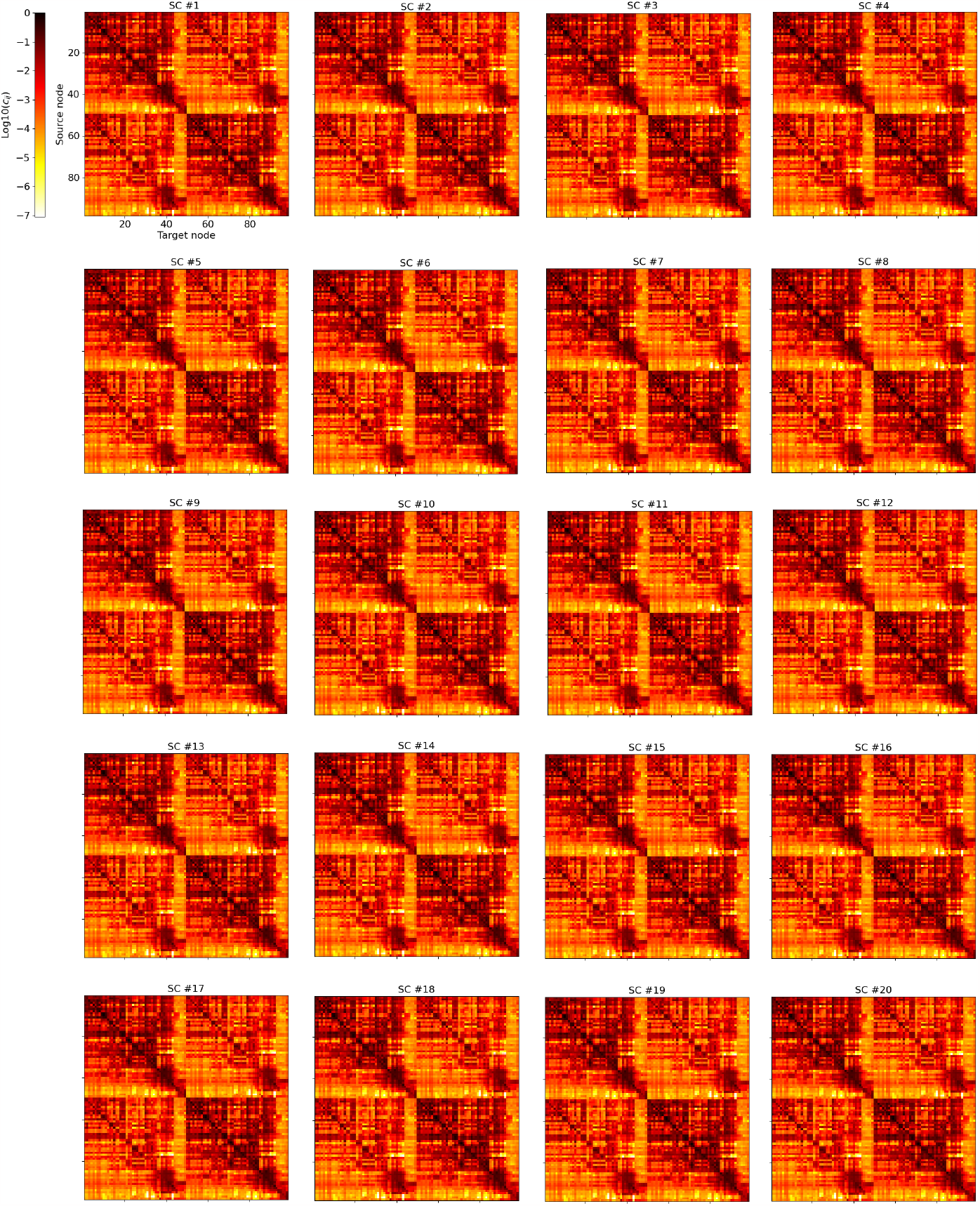
Allen SC matrices used for statistical analysis. For the 20 generated SC, we display the new weight matrix. We show the logarithm of the weight of each connection.

**Figure S2.**
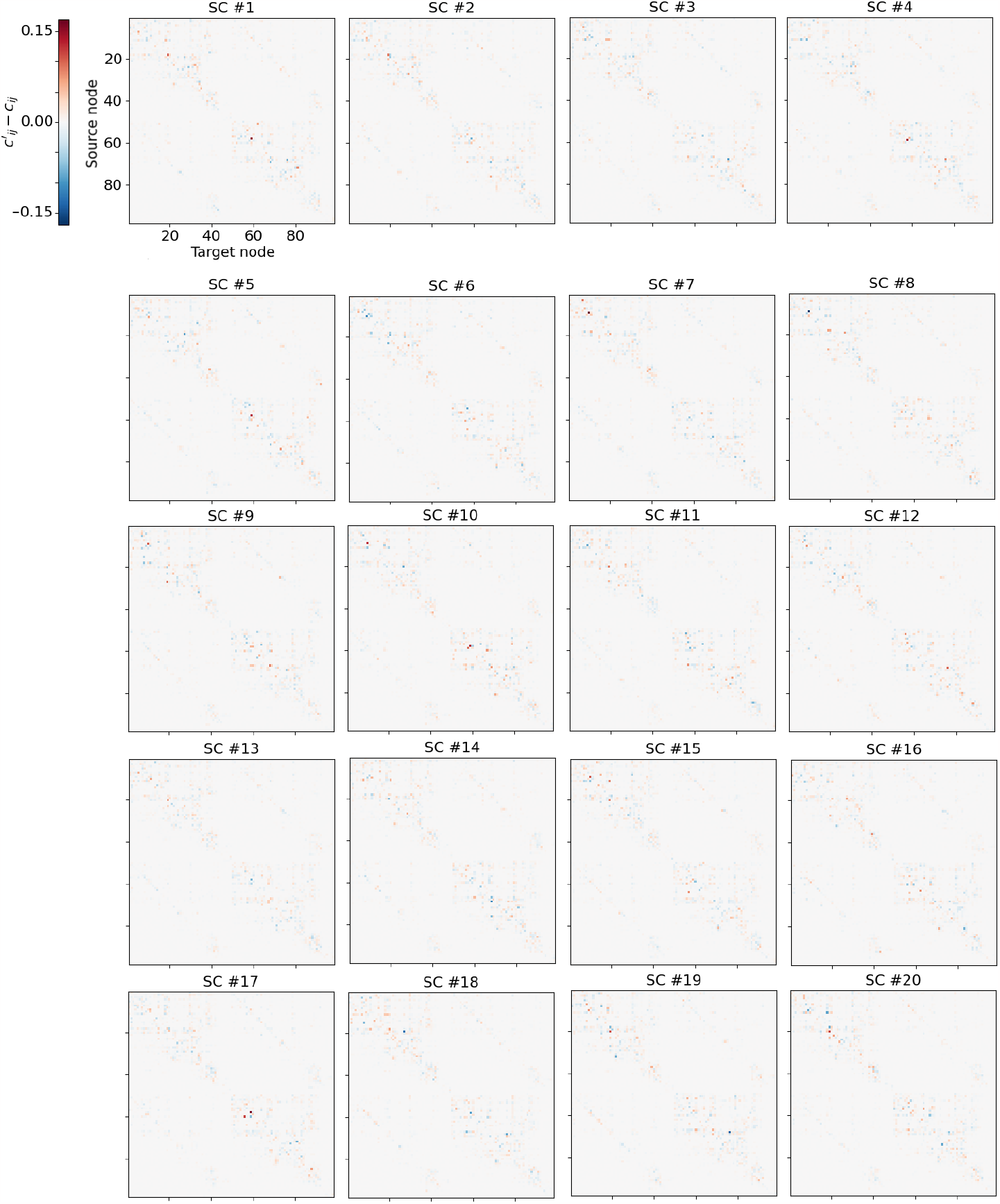
Weight value difference between the additional SC matrices and the original Allen connectome. For the 20 generated SC 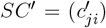, we display the weight difference 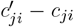 between each respective matrix entry value.

**Figure S3.**
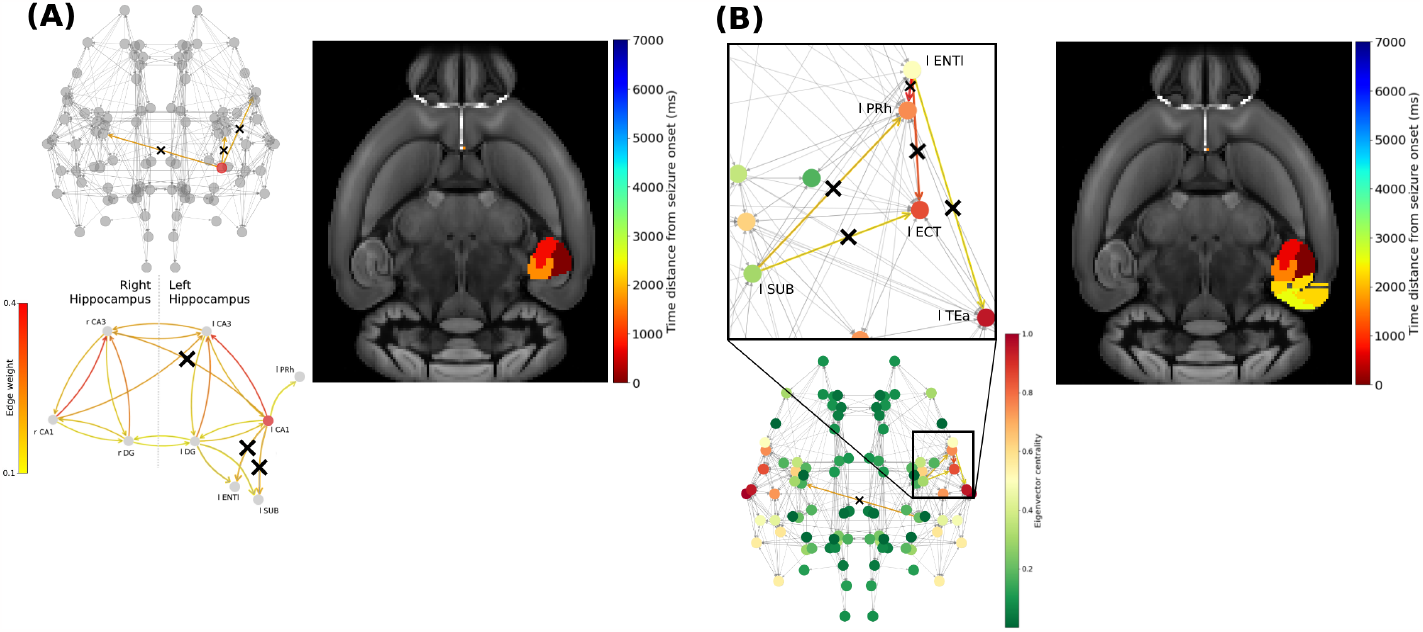
Widespread seizure prevention by edge removal in the left-field CA1.. The upper left panels show the brain network. Crosses indicate the removed connections, and the edges’ color shows their respective weight. The right panels show the time distance between seizure initiation in the EZ and its onset in each brain area. **(A)** Seizure propagation in the Allen connectome after removing the inter-hemispheric strong connection l CA1 ↛ r CA3, and the two strong connections leading from the EZ to outside of left Hippocampus (namely the lateral part of the Entorhinal Cortex ENTl and the Subiculum (SUB)). The lower left panel illustrates the left extra-hippocampal connections and their weights values are given in the left colorbar. **(B)** Seizure propagation after removing the inter-hemispheric strong connection l CA1 ↛ r CA3, and all strong connections (*c*_*ji*_ *>* 0.1) leading from l ENTl, l SUB to networks hubs, namely the left Ectorhinal Cortex (ECT), Perirhinal Cortex (PRh) and Temporal Association (TEa) areas. The color of the nodes show their normalized eigenvector centrality.

The Functional Connectivity (FC) matrix *FC*_*ij*_ is measured by the Pearson Correlation Coefficient *r*, i.e. by correlating simulated BOLD time series data between every pair of nodes in our network graph (Stephan and Friston, 2009; Ladwig et al., 2022):

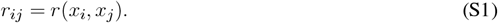

**Figure S4.**
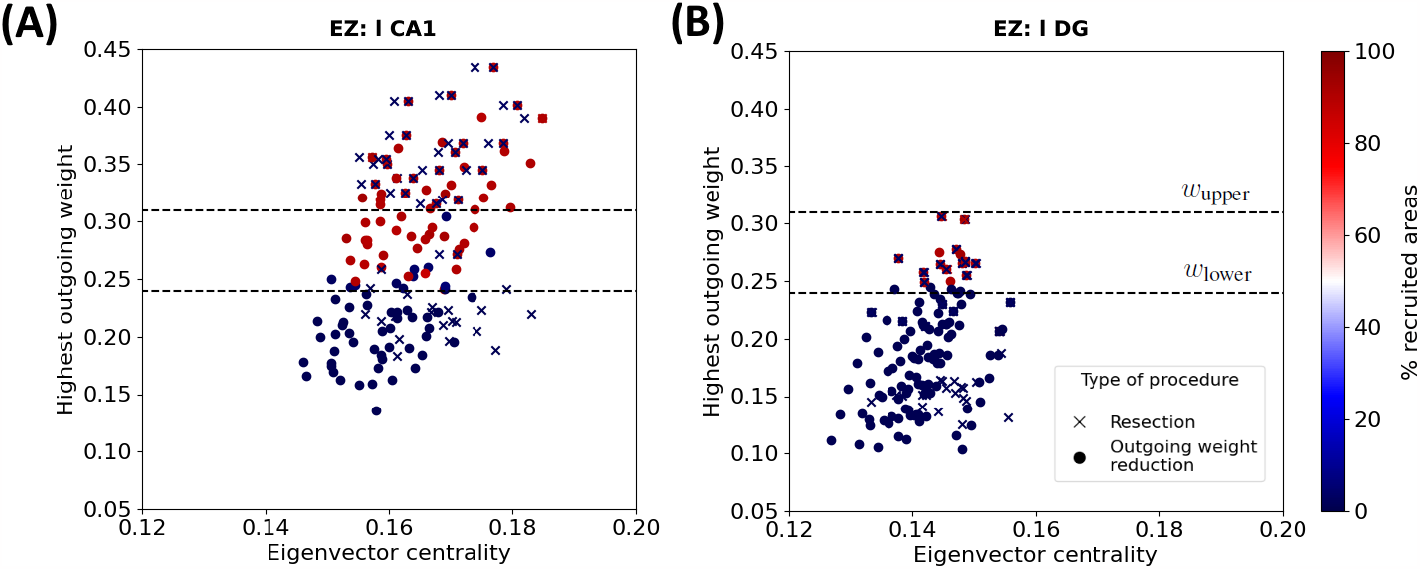
EZ connectivity and percentage of seizure-recruited areas after outgoing weight reduction and edge resection in the 20 generated connectomes. Each point shows the eigenvector centrality and the largest outgoing weight value in one of the 20 randomized connectomes of **(A)** left-field CA1 and **(B)** left Dentate Gyrus, before and after applying outgoing weight reduction (circles) and edge removal strategies presented in **Figure** (6) and **Figure** (7) (crosses). The color of each point shows the percentage of recruited brain areas when the seizure starts in the corresponding node. The two dashed lines show the upper (resp. lower) threshold of the largest outgoing weight from separating the widespread from localized seizures.

**Figure S5.**
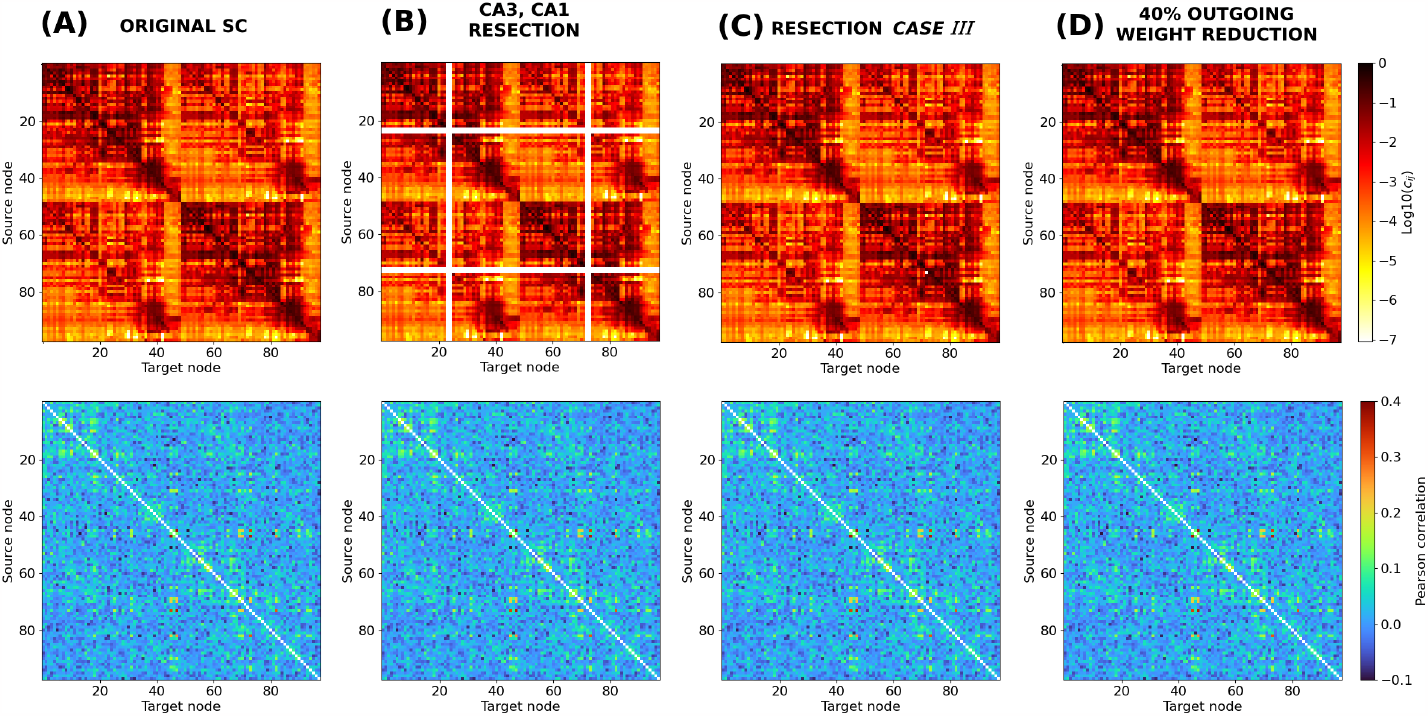
Connectome structure and functionality after resection and outgoing weight reduction procedures. We show the network structural connectivity (top) and functional connectivity (bottom) matrices. **(A)** Original connectome, **(B)** after the resection of left and right CA1 and CA3 areas, following the procedure presented in (Melozzi et al., 2017), **(C)** after applying resection *Case III* and **(D)** after applying l CA1 outgoing weight reduction of 40%.

In **Figure** S5, we show the original Allen SC matrix (panel (A)) together with modified ones, following effective seizure localization strategies. Panel (B) shows the SC after applying a standard resection technique presented in Melozzi et al. (2017). This approach consists in the removal of the CA1 and CA3 areas in both hemispheres. Panel (C) shows the SC after applying resection *Case I*, e.g. the resection of a single connection l CA1 ↛ l CA3, and panel (D) shows the SC after applying a 40% outgoing weight reduction on l CA1. With the latter two options, we minimize the structural modifications compared to the CA1, CA3 resection procedure (Melozzi et al. 2017). By setting our system at a “healthy” (i.e., modeling all network nodes as non EZ), we do not manage to observe notable difference between the FC produced by the original SC matrix and the FC matrices produced with the different intervention procedures introduced in our manuscript. Here we show only a few typical examples. Note that we have also attempted to estimate matrix distances, e.g. by calculating the Pearson correlation coefficient between different pairs of matrices (see e.g. (Popovych et al., 2021; Manos et al., 2023)), but we did not observe striking dissimilarities (results not shown here). It is also worth noting that at this stage we did not aim to compare our simulated time series (or FC matrices) to experimental neuroimaging mice signals.

## Notes

### Competing Interest Statement

The authors have declared no competing interest.

